# STING activation promotes autologous type I interferon-dependent development of type 1 regulatory T cells during malaria

**DOI:** 10.1101/2022.09.04.506109

**Authors:** Yulin Wang, Fabian De Labastida Rivera, Chelsea L. Edwards, Teija C. M. Frame, Jessica A. Engel, Luzia Bukali, Jinrui Na, Susanna S. Ng, Dillon Corvino, Marcela Montes de Oca, Patrick T. Bunn, Megan S. F. Soon, Dean Andrew, Jessica R. Loughland, Fiona H. Amante, Bridget E. Barber, James S. McCarthy, J. Alejandro Lopez, Michelle J. Boyle, Christian R. Engwerda

## Abstract

The development of highly effective malaria vaccines and improving drug treatment protocols to boost anti-parasitic immunity is critical for malaria elimination. However, these efforts are hampered by parasite-specific immunoregulatory networks that are rapidly established following exposure to malaria parasites. Here, we identify stimulator of interferon genes (STING) as a critical mediator of type I interferon production by CD4^+^ T cells during blood-stage *Plasmodium falciparum* infection. STING activation by cyclic guanosine monophosphate-adenosine monophosphate (cGAMP) stimulated *IFNB* gene transcription that promoted development of IL-10 and IFNγ co-producing CD4^+^ T (type I regulatory; Tr1) cells. CD4^+^ T cell sensitivity to STING phosphorylation increased in healthy volunteers following *P. falciparum* infection, particularly in Tr1 cells. Finally, we found the JAK1/2 inhibitor ruxolitinib modulated this innate signalling axis in CD4^+^ T cells to increase parasite-specific Th1 and diminish Tr1 cell responses. These findings identify STING as a critical mediator of Tr1 cell development during malaria.

## Introduction

Malaria is a devastating human disease of global importance. It not only caused an estimated 241 million cases and 627,000 deaths in 2020, but also promoted poverty by imposing health care and socioeconomic costs on communities in malaria endemic areas (WHO, 2021). *Plasmodium falciparum* is responsible for most malaria cases, and despite decades of effort, there is still no licensed vaccine that meets the WHO goal of 75% efficacy against clinical disease (Moorthy et al., 2013). Many factors contribute to this failure including the presence of polymorphic or strain-restricted antigens, sub-optimal vaccine formulations and dosing schedules, as well as imprinted anti-parasitic immune responses that impede rather than enhance vaccine-induced immunity (Montes de Oca et al., 2016c; Nahrendorf et al., 2021; Rogers et al., 2021). In regards to the latter, mounting evidence supports the emergence of potent immune regulatory mechanisms to protect tissues against inflammation following *P. falciparum* infection (Butler et al., 2011; Edwards et al., 2018; Montes de Oca et al., 2016c).

The production of the anti-inflammatory cytokine IL-10 and expression of co-inhibitory receptors by parasite-specific CD4^+^ T cells have been recognised as important components of the immune regulatory networks that arise during malaria (Engwerda et al., 2014; Kumar et al., 2019; Wykes and Lewin, 2018). In fact, IL-10 and IFNγ co-producing CD4^+^ T (type I regulatory; Tr1) cells comprise a substantial fraction of cells responding to parasite antigen stimulation of immune cells from African children living in malaria endemic areas (Boyle et al., 2015; Boyle et al., 2017; Jagannathan et al., 2014; Walther et al., 2009). Evidence from both pre-clinical models (Haque et al., 2010; Haque et al., 2011; Zander et al., 2016) and controlled human malaria infection (CHMI) studies (Montes de Oca et al., 2016c), indicates that type I interferons (IFNs) are important drivers of Tr1 cell development during malaria.

Type I IFNs play diverse roles in host immune responses during infections and cancer (Bogdan et al., 2004; Snell et al., 2017). In a pre-clinical malaria model, type I IFNs were shown to suppress the development of anti-parasitic T follicular helper (Tfh) cell responses, thereby limiting parasite-specific antibody production (Zander et al., 2016). Furthermore, we previously showed that type I IFNs act on dendritic cells to suppress the development of Th1 cells, while promoting *Il10* gene transcription in a model of experimental malaria (Haque et al., 2014). We also demonstrated that in volunteers infected with blood-stage *P. falciparum*, type I IFNs suppressed antigen-specific IFNγ production, while promoting parasite-specific IL-10 production (Montes de Oca et al., 2016c). Although we identified CD4^+^ T cells, along with many other immune cell populations, as important sources of type I IFNs, it isn’t known how type I IFNs are induced during malaria and whether CD4^+^ T cell type I IFN production played any role in the development of anti-parasitic immune responses during malaria. Further, the cellular and molecular signalling pathways involved in Tr1 cell development are unknown. This information is important if we wish to manipulate Tr1 cell development and/or activity to improve anti-parasitic immunity in response to vaccine or drug treatments.

Here, we identified increased expression of *TMEM173* (encoding stimulator of interferon genes (STING)) by Tr1 cells, relative to other CD4^+^ T cell subsets, in volunteers infected with blood-stage *P. falciparum*. Furthermore, we uncovered a role for STING in CD4^+^ T cells type I IFN production and the development of Tr1 cell using primary human cells. These results were verified *in vivo* using a pre-clinical malaria model in mice. Together, our findings identify a critical cell signalling axis in CD4^+^ T cells that drives the development of Tr1 cell during malaria, thus providing a potential means to manipulate this key CD4^+^ T cell subset to improve anti-parasitic immunity either in the context of vaccination or drug treatment.

## Results

### STING is expressed by Tr1 cells from volunteers infected with blood-stage P. falciparum

We recently described a transcriptional signature for human Tr1 cells during CHMI studies that distinguished them from IFNγ-producing CD4^+^ T (Th1) cells (Edwards *et al*., submitted). Further interrogation of this data set (Figure 1A), revealed that *TMEM173* (ENSG00000184584; encoding STING) was upregulated by Tr1 cells, compared with Th1 cells and other CD4^+^ T cells (Figure 1B). Increased *TMEM173* expression was confirmed in validation experiments (Figure 1C). Pathways analysis of Tr1 cell transcriptomic data predicted that IL-10 was indirectly associated with STING, as well as interferon regulatory factor 3 (IRF3), a transcription factor downstream of STING activation and a key driver of *IFNB1* gene transcription (Sato et al., 2000) (Figure 1D). Thus, STING was more highly expressed in Tr1 cells compared with other CD4^+^ T cell subsets and associated with IL-10 and type I IFN production by these cells.

**Figure 1.**
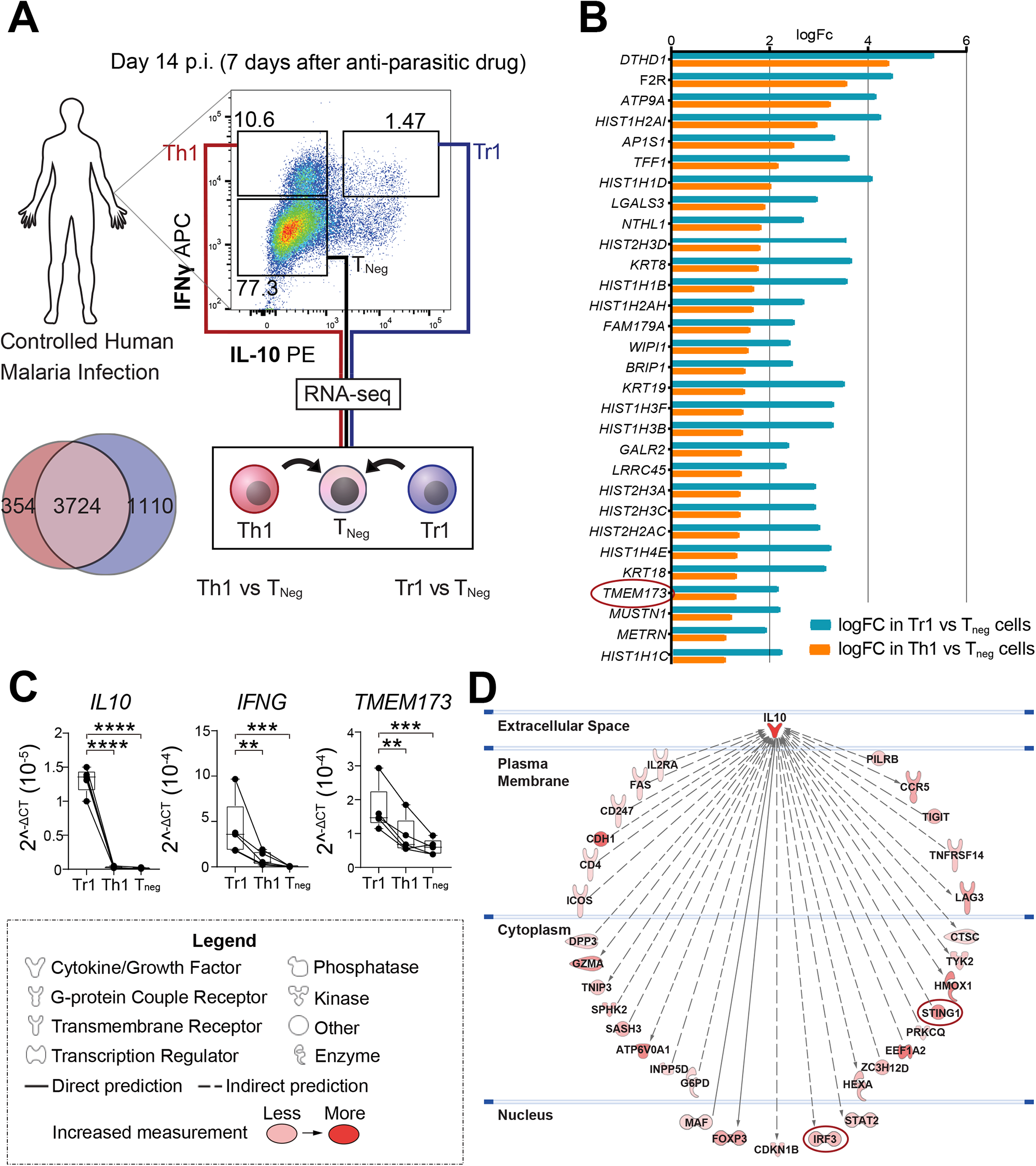
Higher *TMEM173* expression by Tr1 cells compared with Th1 cells during malaria. **A,** A schematic showing the experimental design for the RNAseq analysis of Tr1 and Th1 cells from volunteers participating in controlled human malaria infection (CHMI) studies with *Plasmodium falciparum*. **B,** A list of the top 30 differentially up-regulated genes between Tr1 and Th1 cells from the CHMI study. **C,** Validation of higher *TMEM173* mRNA expression by Tr1 cells, compared to Th1 cells. Human CD4^+^ T cells were isolated from 5 healthy volunteers and then cultured with αCD3ε and αCD28 mAbs plus IL-2 for 3 days. Tr1 and Th1 cells were sorted based on IL-10 and IFNγ expression, as shown in Figure 1A. *TMEM173* mRNA was detected by qPCR and normalized to the housekeeping gene 18S rRNA. Data was Log2-transformed for statistical analysis. Lines connect paired samples and the box shows the extent of the lower and upper quartiles plus the median, while the whiskers indicate the minimum and maximum data points: n=5 samples, paired t-test. Comparisons between Tr1 and other CD4^+^ T cell subsets were made. ** P < 0.01, ***P < 0.001, ****P < 0.0001. **D,** IPA pathways prediction of genes directly or indirectly associated with IL10, as well as the extent of the predicted interaction, as indicated by pink to red colouring.

### Modulation of CD4^+^ T cell STING activation with CRISPR/Cas9 gene editing

To investigate the role of STING in human CD4^+^ T cells, CRISPR/Cas9 gene editing of *TMEM173* was employed. We used a previously reported protocol (Hultquist et al., 2019; Hultquist et al., 2016) to optimise editing STING expression in primary CD4^+^ T cells isolated from peripheral blood of healthy volunteers (Figure S1A-B). CD4^+^ T cells were cultured for 3 days with anti-CD3ε and anti-CD28 monoclonal antibodies (mAbs) in the presence of recombinant IL-2 (Figure 2A), before being edited with a guide RNA (gRNA) targeting the *TMEM173* gene (Figure 2B). Following a further 3 days of cell culture under the same conditions, approximately 73% of cells had excisions in exon 4 of the *TMEM173* gene (Figure S1C-D), reduced *TMEM173* mRNA (Figure 2C) levels and STING protein (Figure 2D). The loss of STING was confirmed by showing stimulation with the STING agonist cGAMP resulting in negligible detection of phosphorylated STING (p-STING) (Figure 2E). Thus, we were able to modify *TMEM173* using CRISPR/Cas9 gene editing, resulting in CD4^+^ T cells that were unable to respond to stimulation with cGAMP.

**Figure 2.**
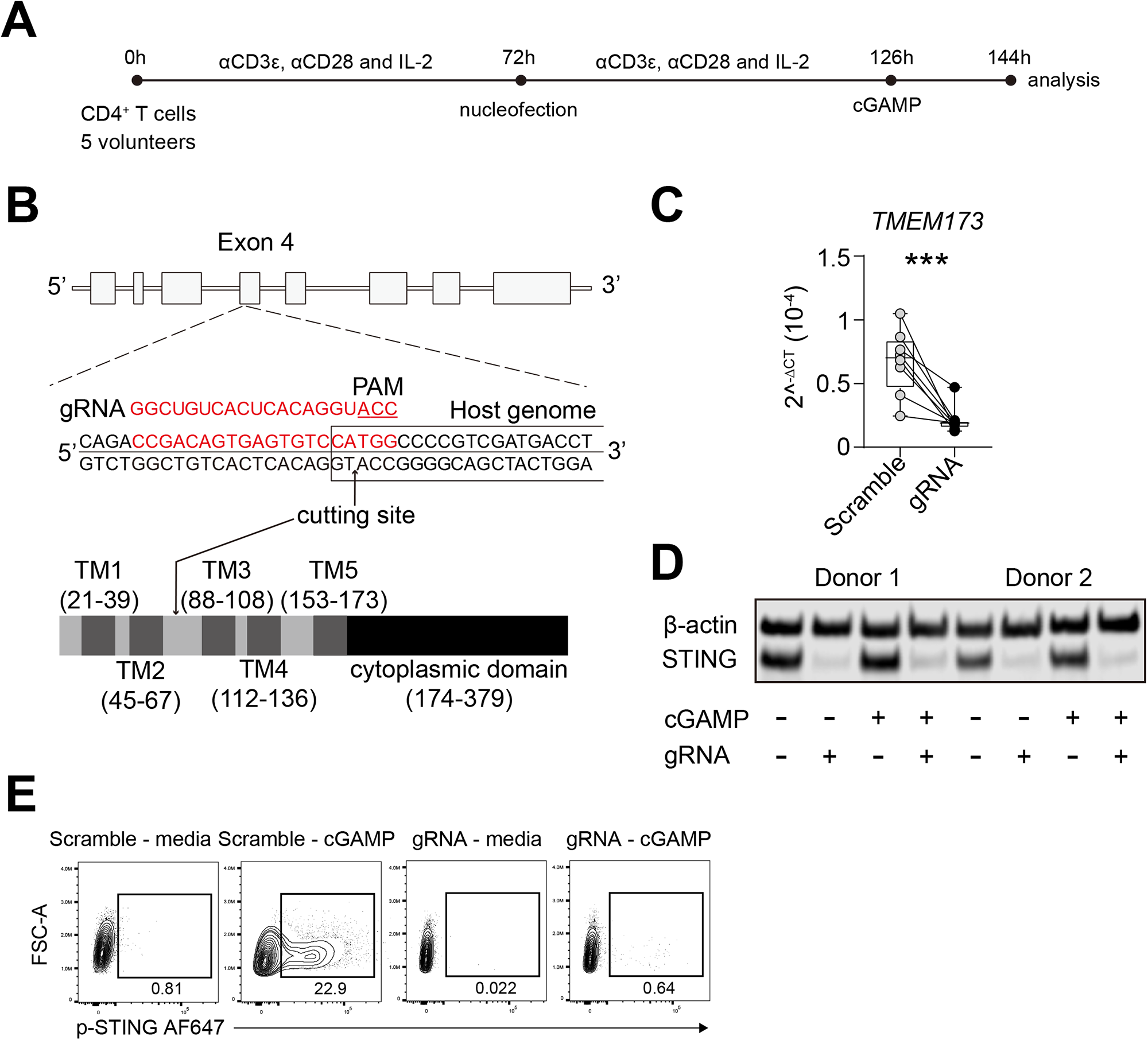
Modulation of CD4^+^ T cell STING expression by CRISPR/Cas9 gene editing. **A,** CD4^+^ T cells from 8 healthy volunteers were stimulated with αCD3ε and αCD28 mAbs plus IL-2 for 72h before nucleofection and then stimulated for another 72h under the same conditions. Cells were treated with or without cGAMP for 18h before analysis. **B,** A diagram showing the gene structure of human *TMEM173* and the CRISPR guide RNA (gRNA) targeting sites within exon 4. The domain structure of the human STING protein showing the 4 transmembrane domains of the N-terminal, responsible for ligand binding and protein dimerization. The C-terminal contains the cyclic dinucleotide (CDN) domain and binding sites for TBK1 and IRF3. **C,** qPCR validation of *TMEM173* mRNA expression in control and CRISPR gRNA treated samples. *TMEM173* mRNA was normalized to 18S rRNA. Data was Log2-transformed for statistical analysis. Lines connect paired samples and the box shows the extent of the lower and upper quartiles plus the median, while the whiskers indicate the minimum and maximum data points: n=8 samples, paired t-test, ***P < 0.001. **D,** Representative western blot of control and CRIPSR gRNA treated samples. β-actin was used as a protein loading control, relative to STING protein levels. **E,** Representative FACS plots showing loss of STING phosphorylation in control and CRIPSR gRNA samples treated, as indicated.

### CD4^+^ T cell STING is required for Tr1 cell development

To identify Tr1 cells without the need for stimulation with strong mitogens such as phorbol myristate acetate (PMA) to detect IL-10 and IFNγ, we used LAG3 and CD49b, which have previously been shown to be highly expressed by Tr1 cells (Gagliani et al., 2013; Huang et al., 2018). Tr1 cells identified by LAG3 and CD49b co-expression peaked at day 4 after stimulation of CD4^+^ T cells with anti-CD3ε and anti-CD28 mAbs plus IL-2, and consistent with previous studies (Brockmann et al., 2018; Gagliani et al., 2013), LAG3 and CD49b co-expressing cells produced the highest amounts of IL-10 and IFNγ, as well as their transcripts (Figure S2). The development of LAG3^+^ CD49b^+^ Tr1 cells was induced by cGAMP (Figure 3A), while the frequency of other CD4^+^ T cell subsets, based on chemokine receptor expression, was decreased (Figures S3A-C), showing that cGAMP activation of STING promoted Tr1 cell development. This was supported by the observation that CRISPR/Cas9 gene editing of *TMEM173* in human CD4^+^ T cells, abolished these effects (Figure 3A). To test whether Tr1 cells were the main CD4^+^ T cell subset responding to cGAMP, we examined STING phosphorylation following cGAMP stimulation of activated CD4^+^ T cells. Indeed, the highest frequency of p-STING^+^ cells were found amongst LAG3^+^ CD49b^+^ CD4^+^ T cells (Figure 3B). Stimulation of CD4^+^ T cells with cGAMP also increased STING-dependent transcription of *IL10*, *IFNG* and *IFNB1*, but this was abrogated following *TMEM173* gene editing (Figures 3C-E). Notably, cGAMP stimulation resulted in no significant increase to the transcription of other type I IFN family members (Figure S3D), suggesting selective induction of *IFNB1* amongst the type I IFN family of genes in CD4^+^ T cells following STING activation. Thus, CD4^+^ T cell STING promotes Tr1 cell development and its activation in these cells drives *IL10*, *IFNG* and *IFNB1* transcription.

**Figure 3.**
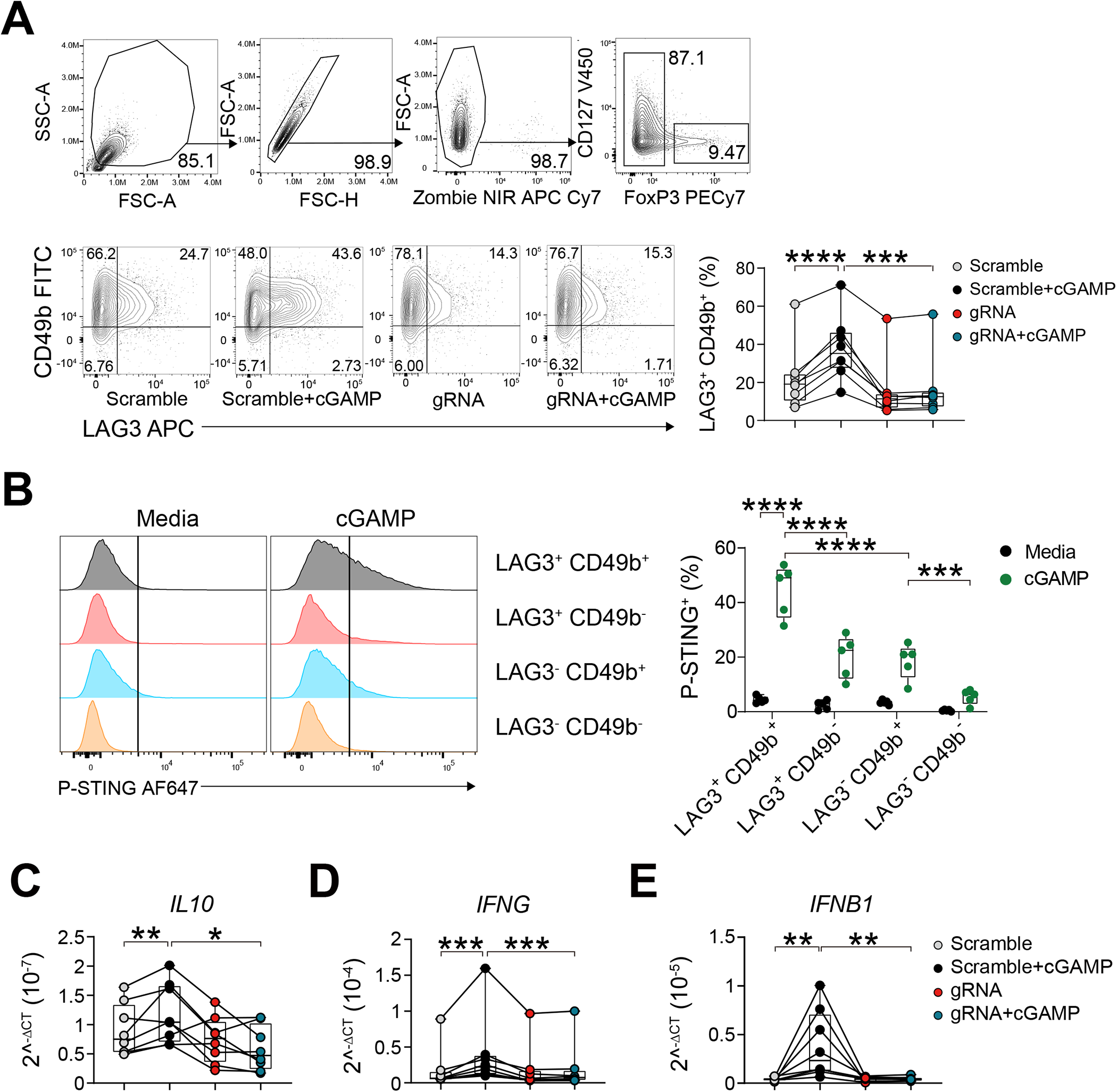
CD4^+^ T cell STING activation promotes Tr1 cell development. Human CD4^+^ T cells were cultured and subjected to CRISPR/Cas9 *TMEM173* gene editing as shown in Figure 2. **A,** The gating strategy used to assess changes in human CD4^+^ T cells. Cells were gated on single cells, live cells and conventional CD4^+^ T cells (FoxP3^-^) before further analysis. Representative plots and enumeration showing the frequency of LAG3^+^ CD49b^+^ CD4^+^ T cells following CRISPR-cas9-mediated modification of *TMEM173* expression. Lines connect paired samples and the box shows the extent of the lower and upper quartiles plus the median, while the whiskers indicate the minimum and maximum data points. **B,** Representative histograms and enumeration showing the frequencies of p-STING-positive LAG3^+^ CD49b^+^, LAG3^+^ CD49b^-^, LAG3^-^ CD49b^+^ and LAG3^-^ CD49b^-^ CD4+ T cell subsets. The box shows the extent of the lower and upper quartiles plus the median, while the whiskers indicate the minimum and maximum data points. **C, D, E,** The expression of *IL10*, *IFNG* and *IFNB1* in the control and *TMEM173*-modified cells with and without cGAMP activation was measured by qPCR. Data was Log2-transformed for statistical analysis. Lines connect paired samples and the box shows the extent of the lower and upper quartiles plus the median, while the whiskers indicate the minimum and maximum data points: n=8 (**A, C-E**) or n=5 (**B**), paired t-test. Comparisons were made between scramble vs scramble + cGAMP, scramble + cGAMP vs gRNA + cGAMP, gRNA + cGAMP vs gRNA. *P < 0.05, **P < 0.01, ***P < 0.001, ****P < 0.0001.

### STING-dependent IFNβ1 production by CD4^+^ T cells drives Tr1 cell development

In addition to having potent anti-viral activities, we and others have previously reported that type I IFNs modulate CD4^+^ T cell responses during experimental and clinical malaria, including suppressing Th1 and Tfh cell responses (Haque et al., 2011; Haque et al., 2014; Montes de Oca et al., 2016c; Zander et al., 2016). Given the strong STING-dependent induction of *IFNB1* by Tr1 cells, we next examined whether this type I IFN production was needed for Tr1 cell development and/or maintenance (Figure 4A). Following CD4^+^ T cell activation, as above, we found that STING-dependent expansion of Tr1 cells was abrogated by blocking type I IFN signalling with an antibody directed against the type I IFN receptor (IFNR) (Figure 4B). We also observed reduced *IL10* and *IFNG* transcription when type I IFN signalling was blocked (Figure 4B), as well as diminished *IFNB1* induction following IFNR blockade (Figure 4B), suggesting a positive feedback mechanism for *IFNB1* transcription. To directly link CD4^+^ T cell autologous STING-dependent IFNβ1 production with Tr1 cell development, we tested whether Tr1 cell development from STING-deficient CD4^+^ T cells could be rescued by exogenous IFNβ1 (Figure 4C), and indeed this was the case (Figure 4D). We also found that supplementation of CD4^+^ T cells with IFNβ1 alone induced *IL10* and *IFNG* transcription, as well as *IFNB1*, again supporting a positive feedback loop for transcription of this cytokine (Figure 4D). Together, these results show that STING-dependent IFNβ1 production by CD4^+^ T cells drives Tr1 cell development.

**Figure 4.**
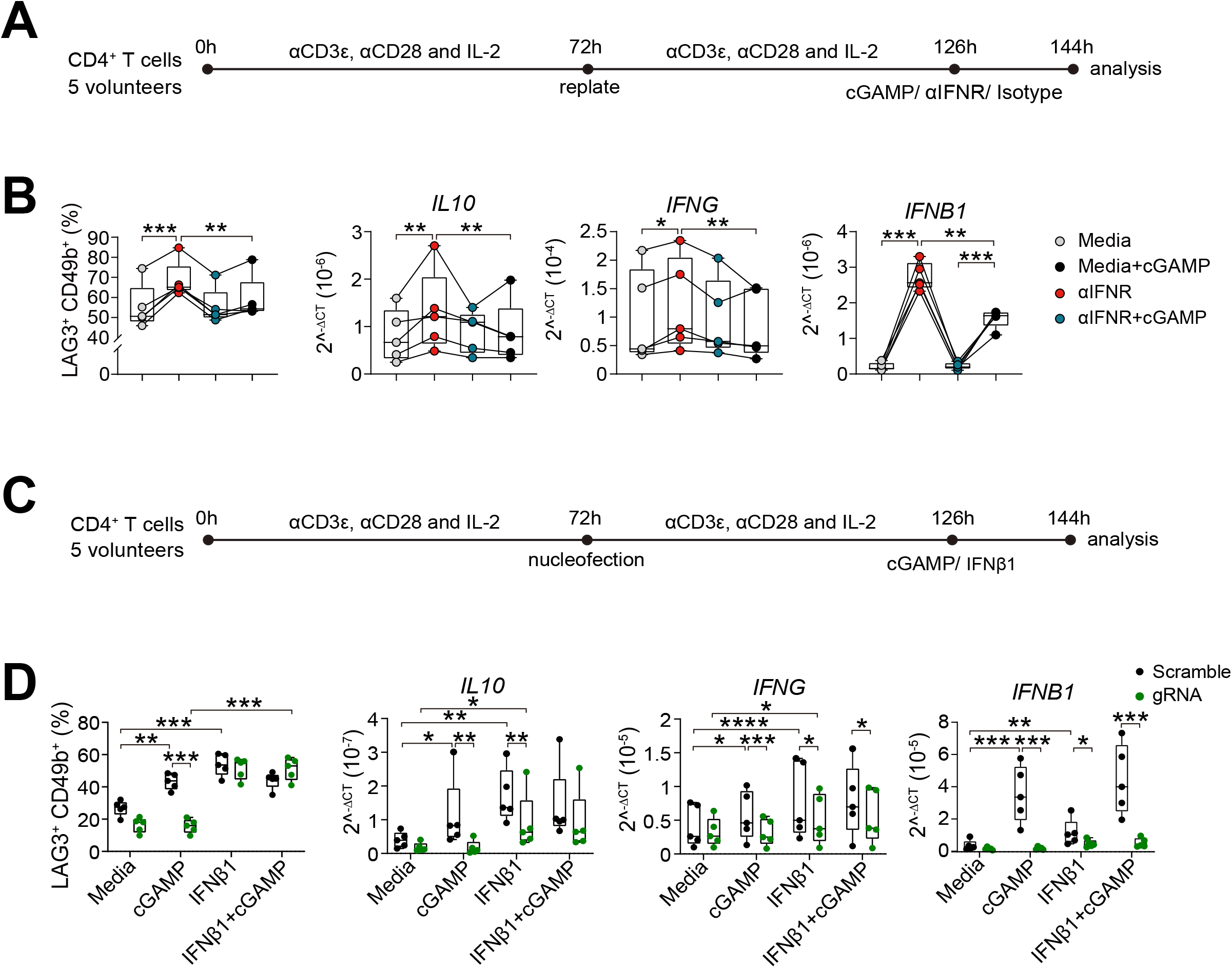
STING-dependent IFNβ1 production by CD4^+^ T cells drives Tr1 cell development. **A**, CD4^+^ T cells were stimulated with αCD3ε and αCD28 mAbs plus IL-2, as shown, prior to treating with an antibody against type I IFN receptor (αIFNR), isotype control mAb or cGAMP for 18h before analysis, as indicated. **B,** Cells were gated on conventional CD4+ T cells (FoxP3^-^), as shown in Figure 3A. The frequency of LAG3^+^ CD49b^+^ CD4^+^ T cells, as well as *IL10*, *IFNG* and *IFNB1* mRNA levels in each treatment group were measured. qPCR data was normalized to the housekeeping gene *18S* rRNA, and was Log2-transformed for statistical analysis. Lines connect paired samples and the box shows the extent of the lower and upper quartiles plus the median, while the whiskers indicate the minimum and maximum data points: n=5, paired t-test. **C,** CRISPR/Cas9 modification of *TMEM173* in CD4^+^ T cells stimulated with αCD3ε and αCD28 mAbs plus IL-2, as shown, prior to treating with 100 ng/μl recombinant IFNβ1 and/or 30 μ g/ml cGAMP 18 hours before analysis, as indicated. **D,** The frequency of LAG3^+^ CD49b^+^ CD4^+^ T cells and *IL10*, *IFNG* and *IFNB1* mRNA were measured in CD4^+^ T cells treated as indicated. Data was Log2-transformed for statistical analysis. The box shows the extent of the lower and upper quartiles plus the median, while the whiskers indicate the minimum and maximum data points n=5, paired t-test. For all statistical tests *P < 0.05, **P < 0.01, ***P < 0.001, ****P < 0.0001.

### CD4^+^ T cell STING is required for Tr1 cell development in experimental malaria

To extend the above studies to an *in vivo* setting, we used an experimental model of severe malaria caused by infection of C57BL/6 mice with *P. berghei* ANKA (*Pb*A). We employed PbTII mice (Enders et al., 2021; Fernandez-Ruiz et al., 2017), a TCR transgenic mouse line that produce CD4^+^ T cells specific for I-A^b^-restricted *PbA* heat shock protein 90 expressed by all rodent and human *Plasmodium* species, and crossed these with *Tmem173*-deficient mice (James et al., 2018; Jin et al., 2011) to generate STING-deficient PbTII cells (PbTII*^ΔSting^*). Wild-type control PbTII cells were generated by crossing PbTII TCR transgenic mice with congenic (CD45.1) C57BL/6 mice to produce mice expressing both *cd45.1* and *cd45.2* alleles (PbTII^WT^). We then isolated PbTII*^ΔSting^*and PbTII^WT^ cells from these animals to test whether CD4^+^ T cell STING was needed for Tr1 cell development *in vivo*. These cells were transferred at an equal mix (10^6^ total) into congenic (CD45.1) C57BL/6 recipient mice the day before *Pb*A infection (Figure 5A). Cell frequencies and cytokine production were subsequently measured at day 4 post-infection (p.i.) in the spleen when Th1 cell responses peak in this tissue in this model (Figure S4). A decrease in the proportion of splenic PbTII*^ΔSting^* cells producing IL-10 and an increase in those producing IFNγ was observed, relative to control PbTII^WT^ cells (Figure 5B; Figure S5A-B). Furthermore, there was a decrease in the proportion of IL-10^+^ IFNγ^+^ PbTII*^ΔSting^* Tr1 cells, compared to control PbTII^WT^ cells (Figure 5C). This was also accompanied by a decreased frequency of PbTII*^ΔSting^*cells producing granzyme (Gzm) B and perforin, relative to PbTII^WT^ cells (Figure S6A-C). The expression of these cytotoxic molecules has previously been associated with both mouse and human Tr1 cells (Chen et al., 2021; Hidalgo et al., 2008). However, differences in Tr1 cells defined by LAG3 and CD49b expression were less consistent between PbTII*^ΔSting^* and PbTII^WT^ cells (Figure 5D). This result suggest that alternative type I IFN cellular sources (not adoptively transferred PbTII*^ΔSting^* cells) were driving expression of LAG3 and CD49b, but not changes in cytokine, GzmB or perforin by CD4^+^ T cells *in vivo*. Hence, these results indicate that CD4^+^ T cell STING promotes IL-10 production while suppressing IFNγ production in a cell intrinsic manner *in vivo* in experimental malaria.

**Figure 5.**
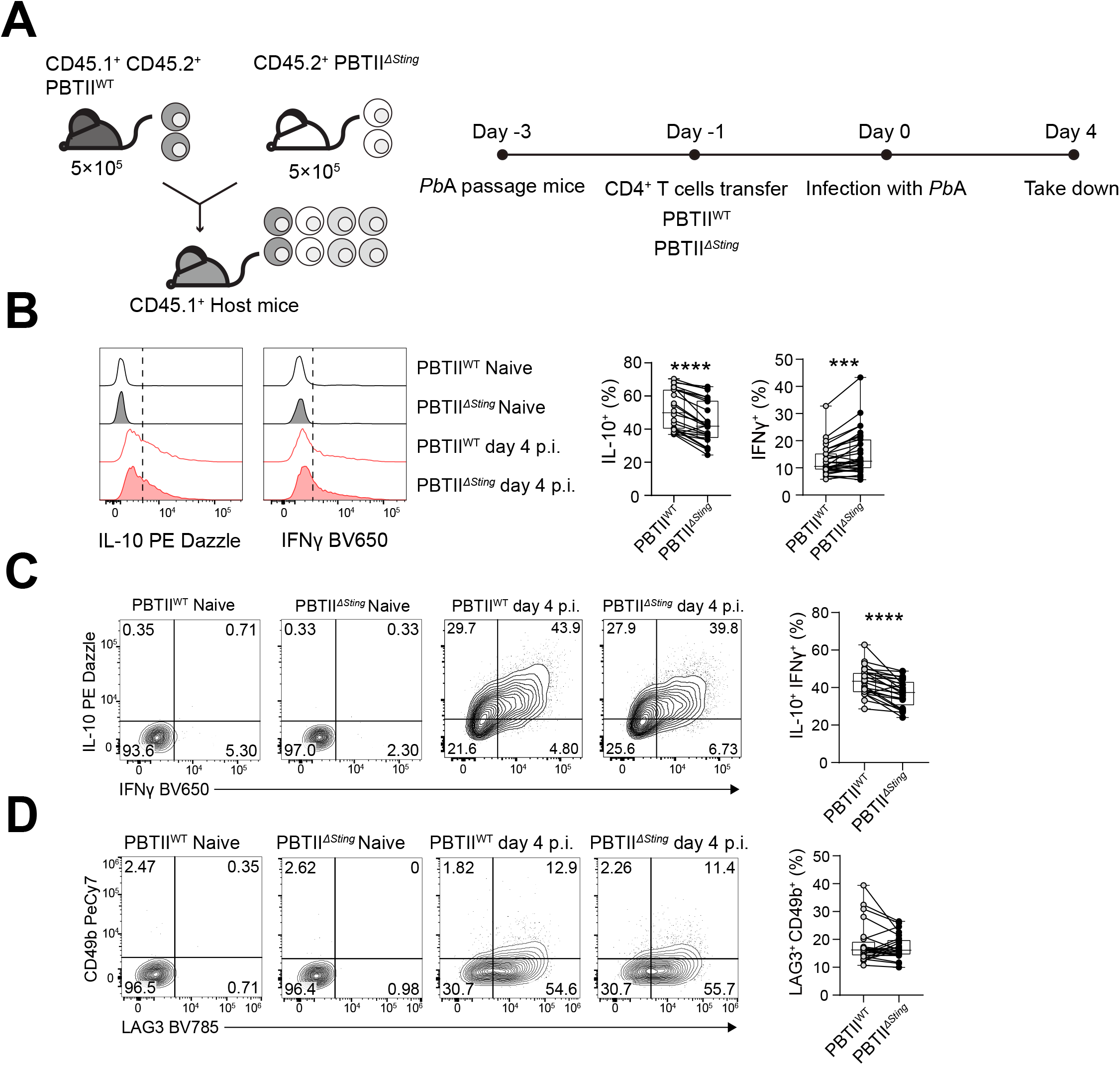
CD4^+^ T cell STING is required for Tr1 cell development in experimental malaria. **A**, 5×10^5^ CD45.2^+^ PbTII*^ΔSting^* and 5×10^5^ CD45.1^+^ CD45.2^+^ PbTII^WT^ cells were transferred into the *Ptprca* (CD45.1^+^) recipient mice at day -1. The mice were infected with *Plasmodium berghei* ANKA (*Pb*A) on day 0 and were assessed on day 4. **B,** Representative histograms and enumeration showing the IL-10- and IFNγ-producing PbTII^WT^ and PbTII*^ΔSting^* cells. **C, D,** Representative plots and enumeration showing the frequencies of IL-10^+^ IFNγ^+^ and LAG3^+^ CD49b^+^ CD4^+^ T cells, respectively. The data in each plot was pooled from 3 independent experiments. Lines connect paired samples and the box shows the extent of the lower and upper quartiles plus the median, while the whiskers indicate the minimum and maximum data points: n = 24, paired t-test, ***P < 0.001, ****P < 0.0001.

### Type I IFN signalling in CD4^+^ T cells drives Tr1 cell development in experimental malaria

We showed that STING-dependent IFNβ1 production by CD4^+^ T cells drives Tr1 cell development of human CD4^+^ T cells *in vitro* (Figure 4). To examine whether cell intrinsic type I IFN signalling was required for Tr1 cell development *in vivo*, we again employed the above model of experimental malaria. However, instead of using PbTII*^ΔSting^* cells, we crossed PbTII mice with *Ifnar*–deficient mice (Hwang et al., 1995; Swann et al., 2007) to generate *Ifnar*-deficient PbTII cells (PbTII*^ΔIfnar^*) that lacked the ability to receive stimulation by type I IFNs. Following transfer of an equal mix (10^6^ total) of PbTII*^ΔIfnar^* and PbTII^WT^ cells into congenic C57BL/6 recipient mice the day before *Pb*A infection, cell frequencies and cytokine production were measured in the spleen at day 4 p.i. (Figure 6A). We found a decreased proportion of splenic PbTII*^ΔIfnar^* cells producing IL-10 or IFNγ, and a decrease in cells producing IFNγ plus IL-10, as well as LAG3^+^ CD49b^+^ Tr1 cells, relative to control PbTII^WT^ cells (Figure 6B-D). As previously observed, the decrease in PbTII*^ΔIfnar^* Tr1 cell frequency, was also associated with a decreased frequency of Tr1 cells producing GzmB and perforin, relative to PbTII^WT^ Tr1 cells (Figure S7). Hence, these results show that CD4^+^ T cell intrinsic type I IFN signalling is required for optimal IL-10 and IFNγ production, as well as Tr1 cell development *in vivo* in experimental malaria. Furthermore, the results suggest that type I IFN signalling plays distinct roles in CD4^+^ T cell IFNγ production, whereby it is needed for induction of IFNγ production, but also promotes the STING-IL-10 axis, that in turn, suppresses IFNγ production by Th1 cells.

**Figure 6.**
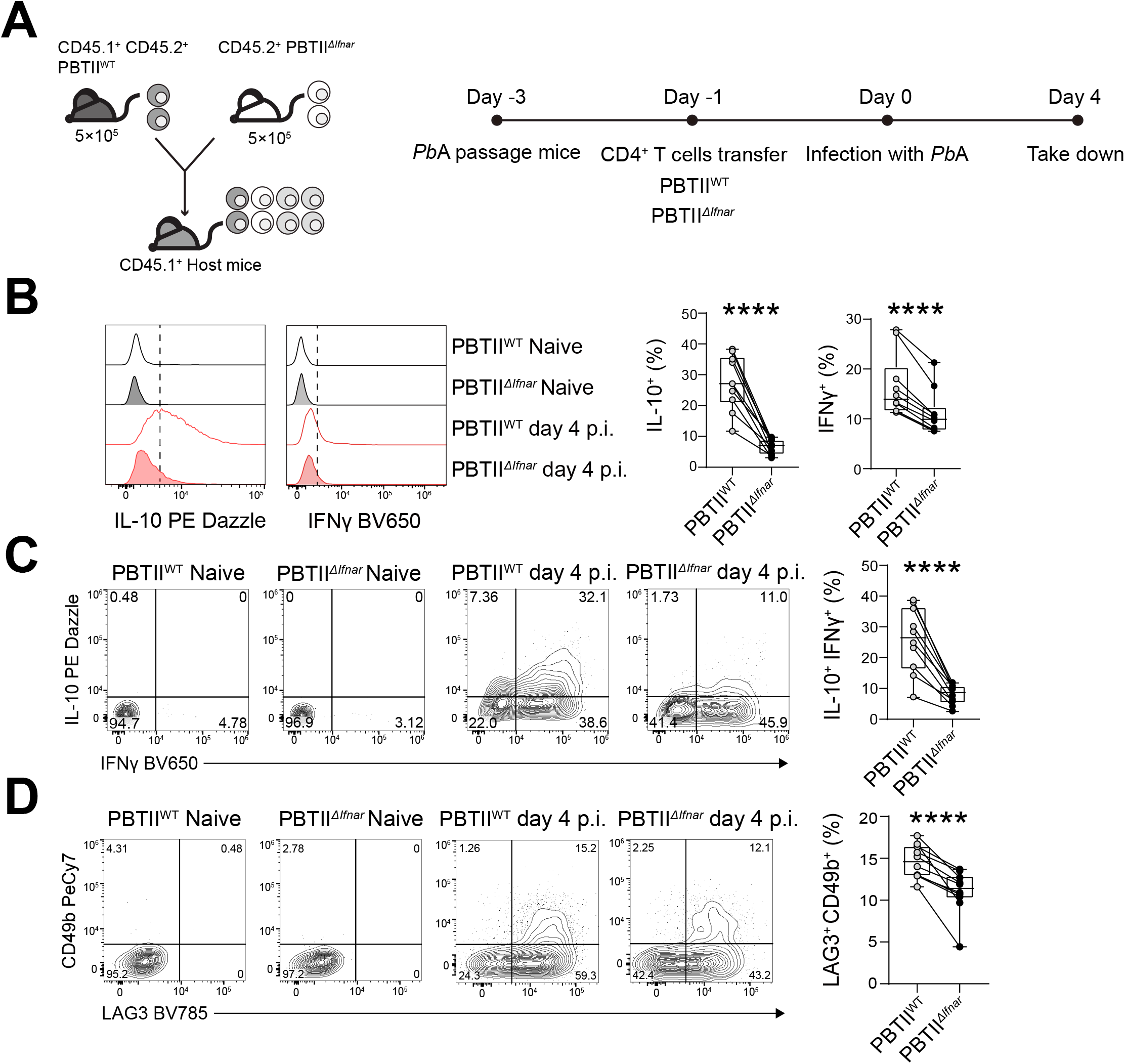
Type I IFN signalling to CD4^+^ T cells drives Tr1 cell development in experimental malaria. **A**, 5×10^5^ CD45.2^+^ PbTII*^ΔIfnar^* and 5×10^5^ CD45.1^+^ CD45.2^+^ PbTII^WT^ cells were transferred into the *Ptprca* (CD45.1^+^) recipient mice at day -1. The mice were infected with *Plasmodium berghei* ANKA (*Pb*A) on day 0 and were assessed on day 4. **B,** Representative histograms and enumeration showing IL-10- and IFNγ-producing in PbTII^WT^ and PbTII*^ΔIfnar^* cells. **C, D,** Representative plots and enumeration showing the frequencies of IL-10^+^ IFNγ^+^ and LAG3^+^ CD49b^+^ CD4^+^ T cells, respectively. The data is pooled from 2 independent experiments. Lines connect paired samples and the box shows the extent of the lower and upper quartiles plus the median, while the whiskers indicate the minimum and maximum data points: n = 10, paired t-test, ****P < 0.0001.

### Tr1 cells from humans infected with P. falciparum are more sensitive to STING activation

Our results above indicate that STING-dependent IFNβ1 production by human CD4^+^ T cells drives Tr1 cell development *in vitro* and a similar STING-dependent pathway promotes Tr1 cell development *in vivo* in experimental malaria. Therefore, we hypothesised that STING would be more readily activated in humans infected with *P. falciparum*. To test this, we examined peripheral blood CD4^+^ T cells from volunteers participating in CHMI studies with *P. falciparum*. We assessed p-STING in CD4^+^ T cell subsets over the course of infection, following stimulation of PBMCs with cGAMP (Figure 7A, Figure S8A). The frequency of cells containing p-STING was heterogeneous among volunteers, but appeared to peak in Tr1 cells in most volunteers at day 15 p.i. (Figure S8B). At this time, the frequency of Tr1 cells expressing p-STING was significantly greater than other CD4^+^ T cell subsets examined (Figure 7B). We next tested whether exposure to parasites increased CD4^+^ T cell sensitivity to STING activation and subsequent development of Tr1 cells. PBMCs collected from CHMI volunteers 15 days p.i. were cultured for 18 hours with uninfected red blood cells (uRBCs) and *P. falciparum*-parasitised red blood cells (pRBCs) with or without cGAMP (Figure 7C). Day 15 p.i. was chosen because this is when anti-parasitic immune responses peak in CHMI volunteers (Montes de Oca et al., 2016c). cGAMP stimulation increased the frequency of Tr1 cells (LAG3^+^ CD49d^+^) at this time point in the presence uRBCs and pRBCs, but this increase was greatest in the presence of pRBCs (Figure 7D).

**Figure 7.**
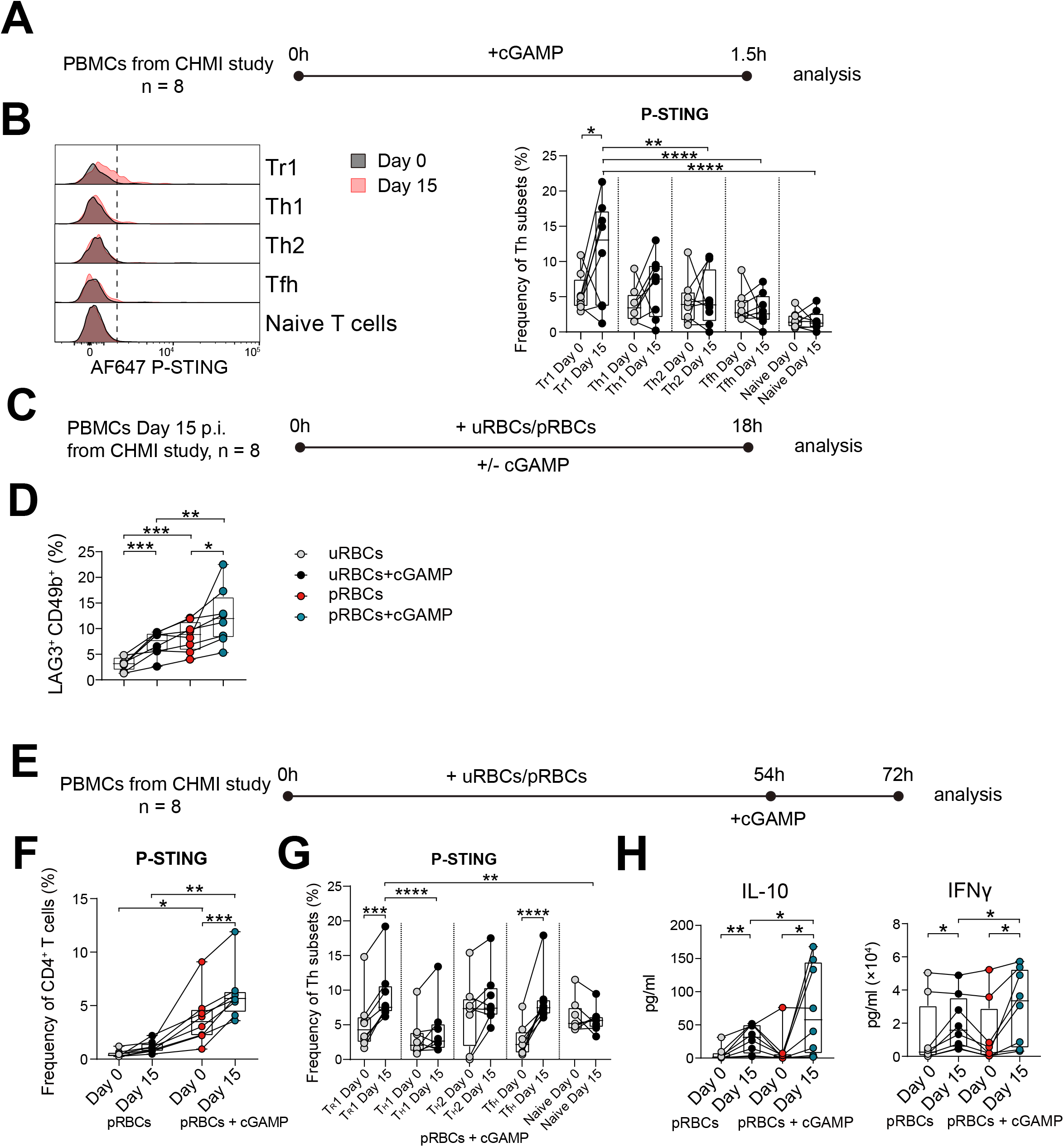
Tr1 cells from humans infected with *Plasmodium falciparum* are more sensitive to STING activation. **A**, PBMCs were isolated from volunteers participating in a controlled human malaria infection study with *P. falciparum* at day 0 and 15 post-infection (p.i.) and stimulated with or without cGAMP for 1.5h before analysis. **B,** Representative histogram and enumeration showing the expression of p-STING in different Th cell subsets day 0 and 15 p.i.. **C,** PBMCs were stimulated with uninfected red blood cells (uRBC) or parasitised RBCs (pRBCs) for 18h, with or without cGAMP before analysis of CD4^+^ T cell subset frequencies. **D,** Tr1 (LAG3^+^ CD49b^+^) cell frequencies as a percentage of CD4^+^ T cells are shown. **E,** PBMCs were stimulated with uRBCs or pRBCs for 72h, and stimulated with cGAMP 18h before analysis. **F,** The frequency of p-STING^+^ CD4^+^ T cells in the presence of pRBCs with or without cGAMP at day 0 and 15 p.i.. **G,** The expression of p-STING in different CD4^+^ T cell subsets in the presence of pRBCs and cGAMP at day 0 and 15 p.i.. **H,** IL-10 and IFNγ produced in the cell culture supernatant in the presence of pRBCs with or without cGAMP at day 0 and 15 p.i.. Lines connect paired samples and the box shows the extent of the lower and upper quartiles plus the median, while the whiskers indicate the minimum and maximum data points (**B, D, F, G** and **H**): n=8, paired t-test. For all statistical tests *P < 0.05, **P < 0.01, ***P < 0.001, ****P < 0.0001.

Finally, we examined how exposure to *P. falciparum* impacted different CD4^+^ T cell subset’s sensitivity to STING activation. We cultured PBMCs from CHMI volunteers taken prior to infection and at day 15 p.i. for 72 hours with uRBCs or pRBCs, with and without cGAMP (Figure 7E). As anticipated from results above, a significantly greater frequency of CD4^+^ T cells from volunteers infected with *P. falciparum* responded to cGAMP activation (Figure 7F), indicating increased sensitivity of CD4^+^ T cells to STING activation following *P. falciparum* infection. Furthermore, a significantly greater frequency of Tr1 (LAG3^+^ CD49b^+^) and Tfh cells (CXCR5^+^ PD1^+^) responded to cGAMP activation following *P. falciparum* infection, but not other CD4^+^ T cell subsets examined (Figure 7G). This increase in Tr1 cell STING sensitivity was associated with a significant increase in antigen-stimulated IL-10 and IFNγ production (Figure 7H). Thus, Tr1 cells in humans infected with *P. falciparum* have increased sensitivity to STING agonists, and this in turn promoted IFNβ-mediated IL-10 production, and subsequently altered the composition of peripheral blood Th cell subsets.

### Ruxolitinib increases the proportion of Th1 cells

After identifying this STING-mediated type I IFN-dependent Tr1 cell development axis, we next attempted to modulate this pathway. Such approaches may be used to overcome malaria vaccine hypo-responsiveness or improve anti-parasitic drug-induced immunity in malaria exposed populations. We recently identified the small molecule JAK1/2 inhibitor ruxolitinib as a potent inhibitor of detrimental type I IFN-dependent, anti-parasitic immune responses in an experimental model of visceral leishmaniasis (VL) in C57BL/6 mice infected with *Leishmania donovani*, as well as in VL patient samples (Kumar et al., 2020). Treatment of human CD4^+^ T cells isolated from healthy volunteers with ruxolitinib at the time of cGAMP stimulation (Figure 8A), limited Tr1 cell development, as well as *IFNB1* mRNA levels (Figure 8B). However, *IL10* and *IFNG* transcription were also suppressed (Figure 8B), indicating a broader impact effect on cytokine gene transcription *in vitro*, not previously observed during *L. donovani* infection *in vivo* (Kumar et al., 2020).

**Figure 8.**
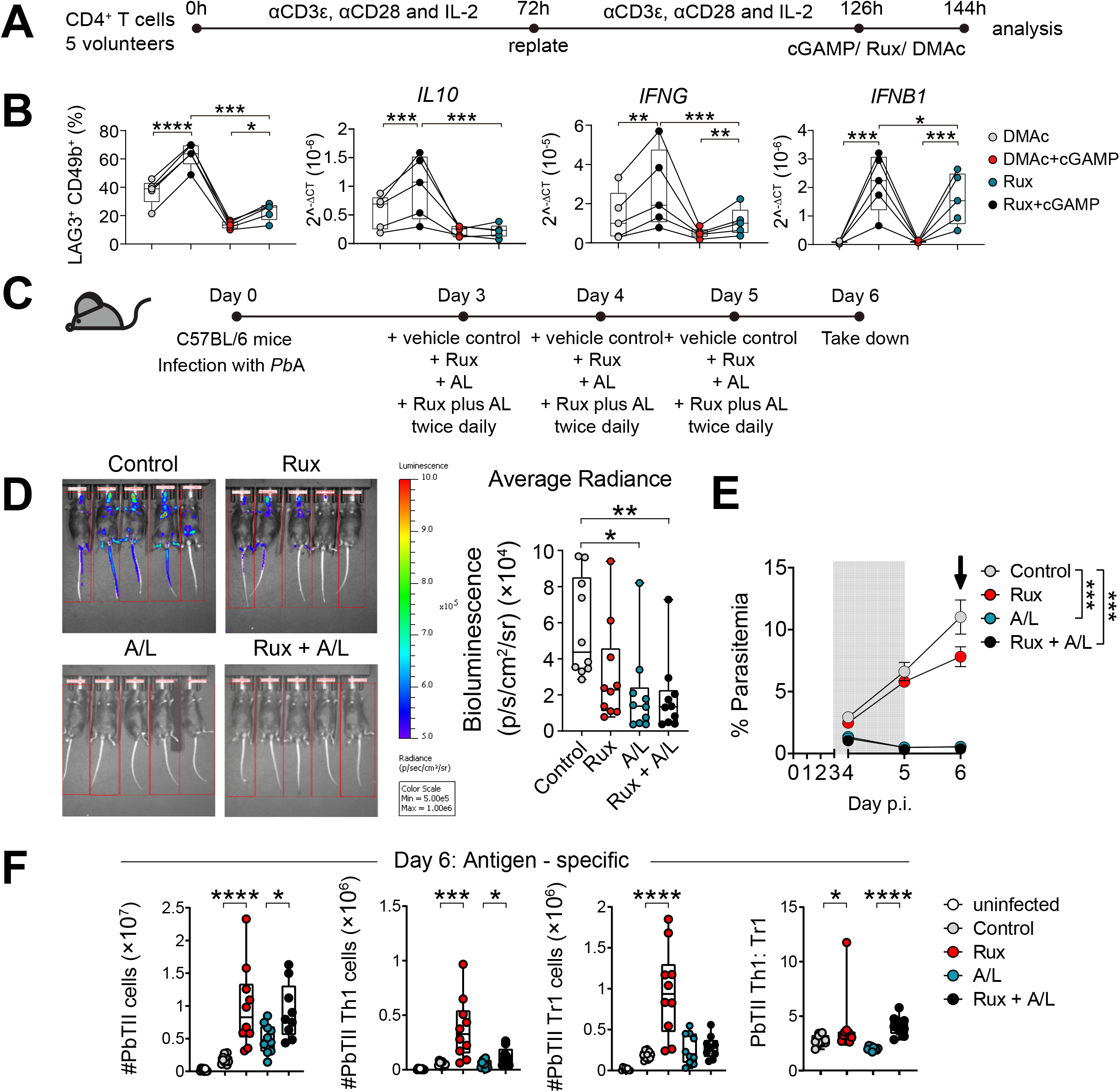
Combination treatment of ruxolitinib and artemether/lumefantrine improves anti-parasitic responses in malaria. **A,** CD4^+^ T cells from the peripheral blood of healthy volunteers were stimulated with αCD3ε and αCD28 mAbs plus IL-2, as shown, prior to treating with Ruxolitinib (Rux) or vehicle (Dimethylacetamide (DMAc)) with or without cGAMP for 18h before analysis, as indicated. **B,** The frequency of LAG3^+^ CD49b^+^ CD4^+^ T cells and *IL10*, *IFNG* and *IFNB1* mRNA were measured in CD4^+^ T cells treated as indicated. Lines connect paired samples and the box shows the extent of the lower and upper quartiles plus the median, while the whiskers indicate the minimum and maximum data points: n=8, paired t-test, *P < 0.05, **P < 0.01, ***P < 0.001, ****P < 0.0001. **C,** C57BL/6J mice were infected with 1×10^5^ luciferase transgenic *Plasmodium berghei* ANKA (*Pb*A-luc) pRBCs intravenously (i.v.) and treated with either DMAC (control), 60mg/kg of Ruxolitinib (Rux), artemether/lumefantrine (A/L; 1.5mg/kg of artemether combined with 9mg/kg of lumefantrine per mouse) or Rux combined with A/L beginning at day 3 post-infection (p.i.) until day 5 post-infection p.i. twice daily and then assessed on day 6 p.i.. **D,** Total body bioluminescence was recorded for 1 minute at day 6 p.i.. **E,** Blood parasitemia was measured daily beginning at day 4 p.i. by flow cytometry. Spleens were harvested for cellular analysis at day 6 p.i. (black arrow). The drug treatment period is depicted by the gray box. **F,** Antigen specific PbTII total, Th1 and Tr1 cell numbers in the spleen at day 6 p.i. are shown for each group, as well as PbTII Th1: Tr1 cell ratios. Boxes show the extent of the lower and upper quartiles plus the median, while the whiskers indicate the minimum and maximum data points (**D, F**). Error bars represent the mean ± SEM (**E**), n= 10 mice per group, data are pooled from two independent experiments (**D, F**), *P < 0.05, **P < 0.01, ***P < 0.001, ****P < 0.0001. **D, F**: Mann-Whitney U test, **E**: One experiment of 2 performed (n = 5/group), Two-way ANOVA with Tukey’s multiple comparisons.

We next tested the *in vivo* utility of for improving anti-parasitic immunity by infecting C57BL/6 mice with *Pb*A and treated them on day 3 p.i. with vehicle controls, ruxolitinib, artemether/lumifantrine (AL) or ruxolitinib plus AL twice daily for 3 days (Figure 8C). This dosing regime was chosen to reflect the AL schedule used to treat human malaria patients (i.e., twice daily for three days) and is currently being tested in CHMI studies (ACTRN 12621000866808). The *Pb*A strain used in these experiments was transgenic for firefly luciferase (Amante et al., 2010; Franke-Fayard et al., 2005), allowing measurements of parasite burden (bioluminescence), as well as blood parasitemia. Mice treated with AL alone or AL combined with ruxolitinib had significantly lower parasite burdens (Figure 8D) and blood parasitemia (Figure 8E), compared to controls, indicating that ruxolitinib wasn’t affecting the anti-parasitic activity of AL, consistent with the lack of drug-drug interaction in a recent Phase I safety trial (Chughlay et al., 2022). Although parasite biomass and blood parasitemia was lower in mice treated with ruxolitinib alone relative to controls, this didn’t reach statistical significance. Interestingly, serum levels of the pro-inflammatory cytokines TNF, IFNγ, MCP-1 and IL-6 were elevated in mice treated with ruxolitinib alone relative to controls, but not in groups treated with AL (Figure S9A). The lower level of serum cytokines in the latter groups likely reflects the rapid control of parasite growth and diminished availability of parasite antigen in these animals after anti-parasitic drug treatment. Although there was no difference in body weights between groups, splenomegaly was more evident in the ruxolitinib alone, AL alone and ruxolitinib plus AL groups relative to controls, possibly reflecting a greater inflammatory response in these animals (Figure S9B).

Finally, we examined the impact of these treatments on antigen-specific CD4^+^ T cells by transferring PbTII^WT^ cells into congenic (CD45.1) C57BL/6 recipient mice the day before *Pb*A infection, treating with ruxolitinib and AL as described above and then assessing PbTII cell phenotype on day 6 p.i.. We found that groups treated with ruxolitinib or ruxolitinib plus AL had higher numbers of total PbTII cells and PbTII Th1 cells, compared to control and AL alone groups, respectively (Figure 8F). When the ratio of Th1:Tr1 cell numbers was calculated, we found a corresponding increase in the ruxolitinib or ruxolitinib plus AL groups, despite the ruxolitinib alone group having significantly higher numbers of PbTII Tr1 cells, relative to controls (Figure 8F). Similar results were observed when we examined splenic polyclonal CD4^+^ T cells in the same animals (Figure S9C), and when cell frequencies were calculated (Figure S9D). Together, these studies show that ruxolitinib can influence host immune responses in experimental malaria either alone or in combination with AL, and support our results from primary human CD4^+^ T cells showing that ruxolitinib can impact the development of Tr1 cells.

## Discussion

In this study, we show that STING is expressed by human Tr1 cells following *P. falciparum* infection. Furthermore, STING can be readily activated by cGAMP to drive CD4^+^ T cell IFNβ1 production that promotes autologous Tr1 cell expansion and/or maintenance, as well as increased IL-10 and IFNγ production following activation. This innate signalling pathway, active in CD4^+^ T cells, could be modulated by the small molecule JAK1/2 inhibitor ruxolitinib when administered with conventional anti-parasitic drugs in experimental malaria, resulting in increased parasite-specific Th1 and diminished Tr1 cell responses.

Tr1 cells have emerged as an important CD4^+^ T cell subset in numerous clinical contexts (Roncarolo et al., 2014; Roncarolo et al., 2018). In malaria, Tr1 cells develop in healthy volunteers infected with *P. falciparum* soon after treatment with anti-parasitic drugs (Montes de Oca et al., 2016c), and were readily detected in African children with malaria (Boyle et al., 2015; Boyle et al., 2017; Jagannathan et al., 2014; Walther et al., 2009). The ability of Tr1 cells to produce IL-10 and express co-inhibitory receptors makes them important for protecting tissues from inflammation (Roncarolo et al., 2014; Roncarolo et al., 2018). However, their immunosuppressive functions may also dampen anti-parasitic immunity, thereby impeding the development of natural, vaccine- or drug-mediated protection against disease (Boyle et al., 2017; Montes de Oca et al., 2016a; Montes de Oca et al., 2016b). Previous studies have shown that the balance between Th1 and Tr1 cell development in mice with experimental malaria is driven by IL-27 (Findlay et al., 2013; Freitas do Rosario et al., 2012; Kimura et al., 2016; Villegas-Mendez et al., 2013) and involves the transcription factors cMaf (Gabrysova et al., 2018) and Blimp-1 (Montes de Oca et al., 2016b). Our findings identify STING expressed by CD4^+^ T cells as another important molecule in Tr1 cell development.

We identified cGAMP-mediated activation of STING as critical for CD4^+^ T cell type I IFN production, and furthermore, showed that this is part of an autologous regulatory loop that suppressed Th1 cell development and promoted Tr1 cell expansion. Of note, only *IFNB* mRNA, and no other members of the type I IFN cytokine family examined, could be consistently measured in human CD4^+^ T cells following activation with cGAMP. Recent studies have shown that in mice, cGAMP can be produced and secreted by cells, then taken up by surrounding cells via the volume-regulated anion channel LRRC8C expressed by T cells to activate STING (Concepcion et al., 2022; Zhou et al., 2020). Hence, one possible mechanism for STING activation in CD4^+^ T cells is via phagocytic cells capturing pRBCs and detecting parasite DNA, as previously described (Gallego-Marin et al., 2018; Sharma et al., 2011), then secreting cGAMP that is taken up by CD4^+^ T cells to activate STING and drive Tr1 cell development during malaria.

In addition to their key roles in innate immune cells, pattern recognition receptors are being increasingly recognised to play important roles in the activation and fate of T cells (Imanishi and Saito, 2020). Previous studies have shown that STING activation in CD4^+^ T cells can induce apoptosis (Larkin et al., 2017; Long et al., 2020) or suppress proliferation (Cerboni et al., 2017). This latter role for STING was shown to be mediated via the inhibition of the metabolic checkpoint kinase, mechanistic target of rapamycin (mTOR), while simultaneously stimulating type I IFN production (Imanishi et al., 2019). Although type I IFNs have previously been shown to induce IL-10 production (McNab et al., 2015; McNab et al., 2014), the cellular and molecular mechanisms responsible have not been fully described. Our findings support a model whereby CD4^+^ T cell IFNβ production in response to STING activation by cGAMP stimulates IL-10 production, as well as Tr1 cell development.

Ruxolitinib is a licensed, orally administered small molecule JAK1/JAK2 inhibitor used to treat polycythemia vera and myelofibrosis (Ajayi et al., 2018). JAK1 and JAK2 phosphorylate signal-transducer and activator of transcription (STAT) molecules which translocate to the nucleus and bind to the promotor of type I IFN-related genes (Seif et al., 2017). Ruxolitinib disrupts the JAK-STAT pathway by competitively inhibiting the ATP-binding catalytic site on JAK1 and JAK2 (Kesarwani et al., 2015). It has been shown in a small case series to improve outcomes in children with type I interferonopathy (Fremond et al., 2016), and to reduce serum type I IFN levels and improve outcomes in adults with refractory dermatomyositis (Ladislau et al., 2018). In another small case study involving STING-associated vasculopathy with onset in infancy (SAVI), ruxolitinib treatment improved respiratory function and skin condition, but also caused recurrence of a respiratory viral infection in one patient (Volpi et al., 2019). We recently showed that ruxolitinib combined with Ambisome, suppressed type I IFN-mediated Tr1 cell IL-10 production, and enhanced Th1 cell-mediated anti-parasitic immune responses in a mouse model of *Leishmania donovani* infection, as well as in whole-blood cells from patients with VL (Kumar et al., 2020). These previous findings, combined with data from this study, as well as a Phase I safety study to examine safety, tolerability, pharmacokinetics, and pharmacodynamics of co-administered ruxolitinib and artemether-lumefantrine in healthy adults have led to a CHMI study with *P. falciparum* to test whether the use of ruxolitinib with AL is safe and can boost anti-parasitic immunity by transiently inhibiting Tr1 cell development and/or functions (ACTRN 12621000866808).

In summary, we have uncovered a Tr1 cell development pathway in mice and humans during malaria that can be targeted by drugs to alter the balance between parasite-specific Th1 and Tr1 cells. These findings have potential applications in strategies designed to improve vaccine efficacy and/or improve anti-parasitic responses following drug treatment, thereby addressing a major bottleneck in efforts to eliminate malaria.

## Acknowledgements

We would like to thank the volunteers who participated in the study, members of the Clinical Malaria laboratory at QIMR and the clinical study team at QPharm (Brisbane, Australia) who conducted the CHMI trial. We thank staff in the QIMR Berghofer flow cytometry laboratory for assistance and staff in the QIMR animal facility for animal husbandry. We thank Nicola Waddell, Ross Loufariotis and Rebecca Johnston for bioinformatics and computational support. This work was made possible through Queensland State Government funding. The research was supported by grants and fellowships from the National Health and Medical Research Council of Australia ((NHMRC; grant numbers 1037304, 1058685, 1132975, 1154265 and 1141632), as well as Australian Post-graduate Awards through Griffith University, School of Natural Sciences and the University of Queensland, School of Medicine. Thanks to Joerg Moehrle and Tim Wells at Medicines for Malaria Venture (MMV) for collaborating on human CHMI studies. Funding for the CHMI platform came for Australia and the UK. We also thank Dr Chris Jeans (QB3 MacroLab, University of California, Berkeley), for supplying purified Cas9-NLS protein for gene editing experiments.

## Author contributions

YW and CRE conceived, planned and wrote the paper. YW, FDLR, CLE, TCM, JAE, LB, JA, SSN, DC, MMO, PTB, MSFS, DA, JRA, FHA, RK and CRE performed experiments and/or analysed data. MSFS, DA, JRL, FHA, BEB, JSM, MJB and JAL provided reagents, assisted with data analysis and/or manuscript editing. SSN and DC analysed the RNAseq data. BEB, JSM and FHA helped plan and execute CHMI studies, as well as edit the manuscript.

## Materials and Methods

### Human and mice ethics

Human ethics approval was provided by the QIMR Berghofer Medical Research Institute Human Ethics Committee (HREC; HREC reference number P1479). Written informed consent was received from all participants. PBMCs were obtained from either healthy volunteers (QIMR Berghofer Medical Research Institute laboratory members) or volunteers participating in a CHMI study with *P. falciparum* (registration number NCT03542149; Table 1)

**Table 1.**
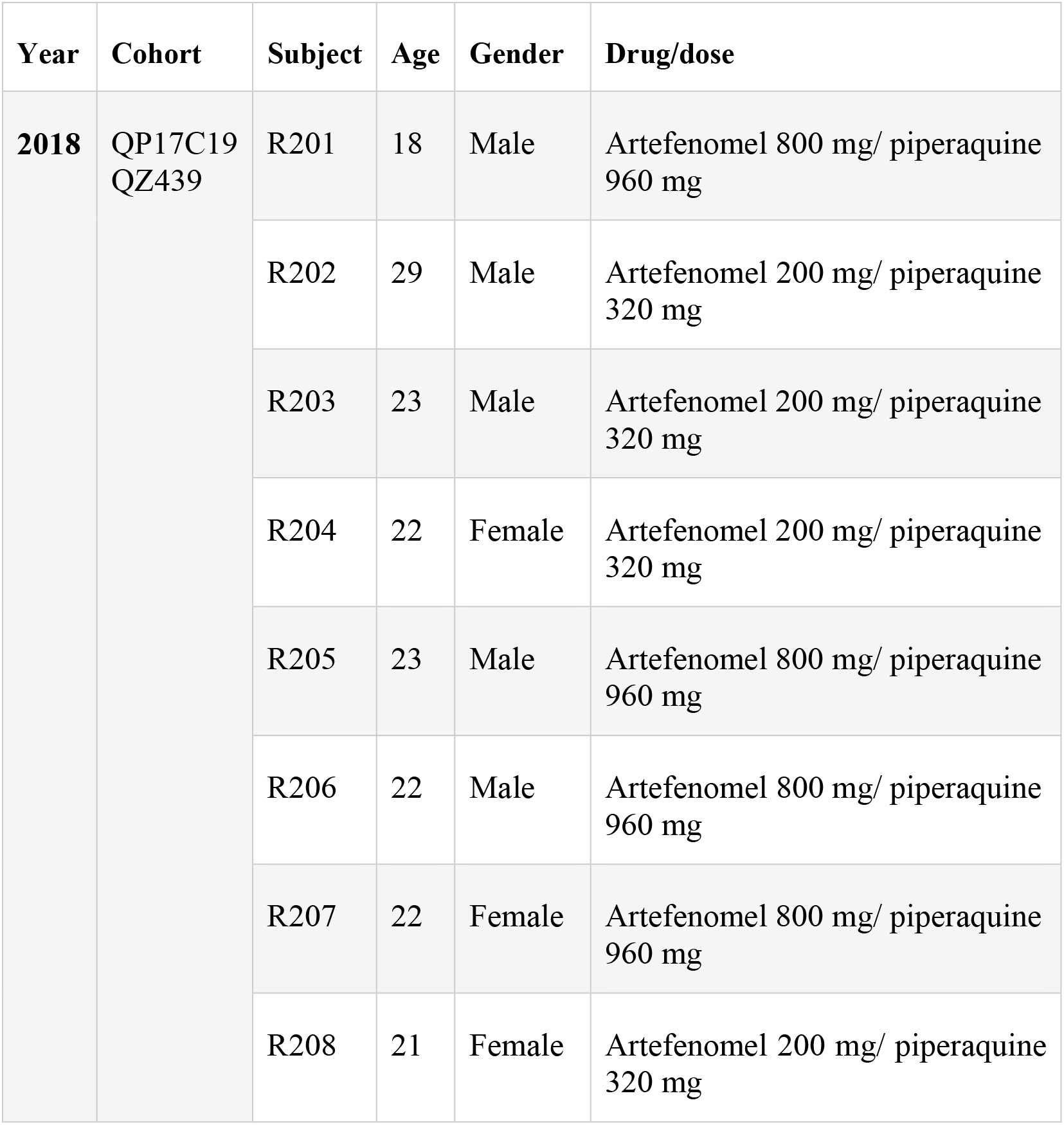
Controlled human malaria infection study participants providing peripheral blood mononuclear cells.

The experimental use of mice followed the “Australian Code of Practice for the Care and Use of Animals for Scientific Purposes” (Australian National Health and Medical Research Council) and was approved by the QIMR Berghofer Medical Research Institute Animal Ethics Committee (Herston, QLD, Australia; approval number: A1707 615M).

### Human primary cells

PBMCs were isolated using Ficoll-Hypaque (GE Healthcare Bio-Sciences AB, Uppsala, Sweden), according to the manufacturer’s instructions. PBMCs were suspended in freezing media (90% (v/v) foetal bovine serum (FBS; Gibco/Thermo Fischer Scientific, Waltham, MA) and 10% (v/v) DMSO (Sigma Aldrich, St Louis, MO)) and stored at -80 °C. PBMCs were thawed and washed with RPMI 1640 Media (Life Technologies, Carlsbad, CA), and rested in complete media (10% (v/v) foetal calf serum (FCS), 100 U/ml penicillin, 100 μg/ml streptomycin (penicillin-streptomycin), 1× GlutaMAX, 1× non-essential amino acids, 1 mM sodium pyruvate, 5 mM HEPES (Gibco) and 0.05 mM 2-mercaptoethanol (Sigma Aldrich), in RPMI 1640 containing l-glutamine (Gibco)) for 40 minutes before any further manipulation. Human CD4^+^ T cells were negatively selected from PBMCs using an EasySep^tm^ human CD4^+^ T cell enrichment kit, according to the manufacture’s protocol (STEMCELL Technologies, Vancouver, Canada).

### Nucleofection

Plates (48- or 96-well, flat bottom) were coated with 10 μg/ml αCD3ε mAb (BioLegend, San Diego, CA) and incubated at 37°C for 5 hours, then stored at 4 °C overnight. Human CD4^+^ T cells were stimulated with plate-bound αCD3ε mAb, 5 μg/ml soluble αCD28 mAb (BioLegend) plus 200 U/ml IL-2 (Miltenyi Biotec, Bergisch Gladback, Germany) for 3 days. 225 μl P3 buffer (Amaxa^tm^ P3 primary cell 96-well Nucleofector^tm^ kit, Lonza, Cologne, Germany) and 50 μl of supplement (Lonza) were mixed to make 275 μl nucleofection buffer. CD4^+^ T cells were then washed with PBS, suspended in P3 buffer (Lonza) to achieve a final concentration of 5×10^5^ to 1×10^6^ cells per 20 μl. 2.4 μl CRISPR guide RNA (gRNA; key resource table) (100 μM) and 2 μl Cas9-NLS (80 μM) (Lonza) were mixed and incubated for 40 minutes at 37 °C. After incubation, 4.4 μl gRNA-Cas9 complex was added to a 20 μl cell suspension in P3 buffer and transferred into a nucleovette (Lonza), placed on Amaxa Nucleofector and 96 well Shuttle (Lonza) with the program EH115. After electroporation, cells were rested in warm media for 20 minutes at 37 °C and stimulated with 25 μl/ml Immunocult (STEMCELL) and 200 U/ml IL-2 for another 3 days.

### Mice

C57BL/6/J (WT) mice were purchased from the Walter and Eliza Hall Institute (WEHI; Kew, VIC, Australia). B6.*Sting*^-/-^ mice (Jin et al., 2011) were kindly provided by Rachel Kuns (QIMR Berghofer Medical Research Institute). Transgenic PbTII mice (Enders et al., 2021; Fernandez-Ruiz et al., 2017) were crossed to B6.*Ptprca* (CD45.1^+^) mice to generate PbTII × B6.*Cd45.1* (PbTII^WT^; CD45.1^+^ CD45.2^+^). B6.*Sting*^-/-^ mice were crossed with PbTII mice to generate *Sting*-deficient PbTII mice (PbTII^Δ*Sting*^; CD45.1^-^ CD45.2^+^). PbTII mice were also crossed to B6.*Ifnar*^-/-^ mice to generate *Ifnar*-deficient PbTII mice (PbTII^Δ*Ifnar*^; CD45.1^-^ CD45.2^+^). All mice were housed under pathogen-free conditions at the QIMR Berghofer Medical Research Institute Animal Facility (Herston, QLD, Australia).

*P. berghei* ANKA infections were established from parasites passaged in C57BL/6J mice. 200 μl of transgenic *P. berghei* ANKA (231c11) parasites (in-house laboratory stock, frozen at -80 °C) expressing luciferase and GFP (Franke-Fayard et al., 2005) were thawed at room temperature and injected via the intraperitoneal route into a passage mouse. 3 days post-infection (p.i.), one drop of blood was collected into 250 μl RPMI/PS with 1 IU/ml heparin from passage mice. Then, 50 μl of this blood suspension was stained with 10 μg/mL Hoechst 33342 (Sigma Aldrich) and 5 μM SYTO 84 (Sigma Aldrich) in RPMI/PS for 22 min at room temperature. Next, 300 μl of RPMI/PS was added, and each sample was acquired on a BD LSRFortessa (BD Biosciences, Franklin Lakes, NJ). The pRBCs were identified as Hoechst 33342^+^ SYTO 84^+^. The passage mouse was sacrificed at >1 % pRBC on day 3 p.i.. Blood was collected from the passage mouse by cardiac puncture, into RPMI/PS containing 1 IU/ml heparin, and centrifuged at 290 g for 7 min at room temperature. RBCs were counted on a haemocytometer (Pacific Laboratory Products, Blackburn, Australia). A parasite inoculum containing 5×10^5^ pRBCs per ml was prepared and mice were injected with 200 μl of the inoculum (1×10^5^ pRBCs) intravenously via the lateral tail vein. Blood parasitemia was measured as described above, while parasite biomass was calculated by measuring luciferase transgenic parasites in live mice, as previously performed (Amante et al., 2010).

### Drug treatment of C57BL/6 mice infected with Plasmodium berghei ANKA

Ruxolitinib (Rux; Chemie Tek, Indianapolis, IN) was diluted in N, N – Dimethylacetamide (DMAC) (Sigma Aldrich) and prepared to 60 mg/kg per 200 μL (diluted in DMAC). Riamet 20/120 (20 mg artemether and 120 mg lumefantrine; AL) (Novartis Basel, Switzerland) was dissolved in DMAC and prepared to 1.5 mg/kg artemether and 9 mg/kg lumefantrine per 200 μL (diluted in DMAC). Mice were administered 200 μL of drugs, as indicated in figure legends or control (DMAC) via oral gavage twice daily on days 3-5 p.i..

### Preparation of splenic single-cell suspension

Mouse spleens were collected and placed in 1% (v/v) FCS in PBS (1% FCS/PBS). Spleens were then mechanically processed through a 100 μm EASYstrainer cell strainer (Greiner Bio-One, Kremsmuenster, Austria) using the back of a 5 ml syringe plunger (Terumo Medical, Tokyo, Japan). Cells were resuspended in 1% FCS/PBS and centrifuged at 350 g, before being lysed with 1 ml Red Blood Cell Lysing Buffer Hybrid-Max (Sigma Aldrich) for 5 minutes at room temperature. Cells were then washed with 10 ml of 1% FCS/PBS and resuspended in 5 ml 1% FCS/PBS at stored at 4°C until required.

### Co-transfer of PbTII cells into recipient mice

Splenic CD4^+^ T cells were isolated by MACS using the mouse CD4^+^ T cell isolation kit (Miltenyi Biotec) according to the manufacturer’s instructions. 100 μl of a single cell suspension was stained with 1 μg/ml CD4 BUV395, TCRβ BUV737, CD45.1 FITC and CD45.2 BV711 (key resource table) to check cell purity (>90%). Then PbTII^Δ*Sting*^ or PbTII^Δ*Ifnar*^ cells were mixed with PbTII^WT^ cells at 1:1 ratio and diluted to 5×10^6^ cells/ml in RPMI/PS. A 200 μl cell suspension (containing 10^6^ cells) was injected intravenously into B6.*Ptprca* (CD45.1^+^) recipient mice.

### Flow cytometry

Human cells: Flow cytometry staining was performed in Falcon 96-Well Clear Round Bottom Tissue Culture (TC)-Treated Cell Culture Microplates (Corning Inc., Corning, NY). Cells were washed and incubated with LIVE/DEAD Fixable Blue (Life Technologies), monocytes blocker (Invitrogen, Waltham, MA) and FcR (Fc receptor) true block (Invitrogen) for 12 minutes at 37°C. Cells were then washed at incubated with fluorescently-conjugated surface staining antibodies (key resource table) for 30 minutes at 37°C. Cells were washed and fixed with BD Cytofix Fixation Buffer (BD Biosciences). After fixation, cells were washed twice and stained with fluorescently-conjugated intracellular staining antibodies (Table 3) for 40 minutes at room temperature. Cells were then washed twice and resuspended in 200 μl PBS at 4°C, before being acquired on a 5-laser Cytek Aurora using SpectroFlo software version 2.3 (Cytek Biosciences, Fremont, CA), then analysed on FlowJo version 10.7.1 (BD Biosciences).

### Pre-stain and stimulation of mouse Tr1 cells

Mouse splenocytes were stained with 4 μg/ml LAG3 BV785 (BioLegend) and 4 μg/ml CD49b PeCy7 (BioLegend) in 30 μl PBS for 30 minutes at 37°C. After staining, 70 μl of complete media was added (described above) along with 100 μl of complete media with 2 x monensin, 25 ng/ml PMA and 1.33 nM ionomycin to the cell suspension and incubated for 3 hours at 37°C.

### Surface and intracellular staining

Mouse pre-stained cells or spleen cells were washed once and incubated with TruStain FcR (BioLegend), True-Stain Monocyte Blocker (BioLegend) and LIVE/DEAD fixable aqua dead cell stain (Life Technologies) for 15 minutes at 37°C. After incubation, cells were stained and fixed following the same procedures as the human sample processing described above.

### PCR for detection of genetic modifications

DNA from CRISPR-edited and control cells were extracted by QuickExtract DNA extraction solution, according to the manufacturer’s instructions (Lucigen, Middleton, WI). The DNA was amplified with *TMEM173* primers (key resource table). 2 μl of genomic DNA (1:3 diluted), 5 μl 5x PCR buffer, 2 μl 25 mM MgCl_2_, 0.5 μl 10 mM dNTP, 0.5 μl 10u M forward primer, 0.5 μl 10 μM Reverse primer and 0.125 U/μl GoTaq DNA Polymerase (Promega, Madison, WI) were mixed to make a 25 μl PCR reaction mix (key resource table). PCR was then performed in a T100 Thermal Cycler (Bio-Rad, Hercules, CA) using conditions outlined in Table 2.

**Table 2.** PCR conditions employed to measure *TMEM173* mRNA

| Step # | Temp °C | Time | Note |
| --- | --- | --- | --- |
| 1 | 94 | 4 min | - |
| 2 | 94 | 30 sec | - |
| 3 | 60 | 30 sec |  |
| 4 | 72 | 1 min | repeat steps 2-4 for 35 cycles |
| 5 | 72 | 2 min |  |
| 6 | 12 | - | hold |

### T7 endonuclease I mismatch assay

After PCR amplification, 10 μl PCR products were incubated with 1.5 μl 10X NEBuffer (New England Biolabs, Ipswich, MA) and 1.5μl Nuclease-free water with the thermal cycler setting: 10 minutes. 95°C; 95–85°C (ramp rate of –2°C/second); 85–25°C (ramp rate of -0.3°C/second). Then 13 μl PCR heteroduplexes were digested by 2 μl of 1 U/μL T7 Endonuclease I (New England Biolabs) at 37°C for 60 minutes. The digestion of CRISPR-edited DNA was visualized by running on a 5 % (w/v) agarose gel.

### Big Dye sequencing

PCR products (1-2 ng per 100 base pairs) were treated with ExoSAP-IT PCR Product Cleanup Reagent (Life Technologies) at a ratio of 5:2 at 37°C for 4 minutes, and 80°C for 1 minute. The PCR products were mixed with 6 pmol *TMEM173* forward primer (Table 3) to make a final volume of up to 10 μl. Then the PCR products were submitted to the QIMR DNA sequencing facility for Big Dye sequencing.

**Table 3.**
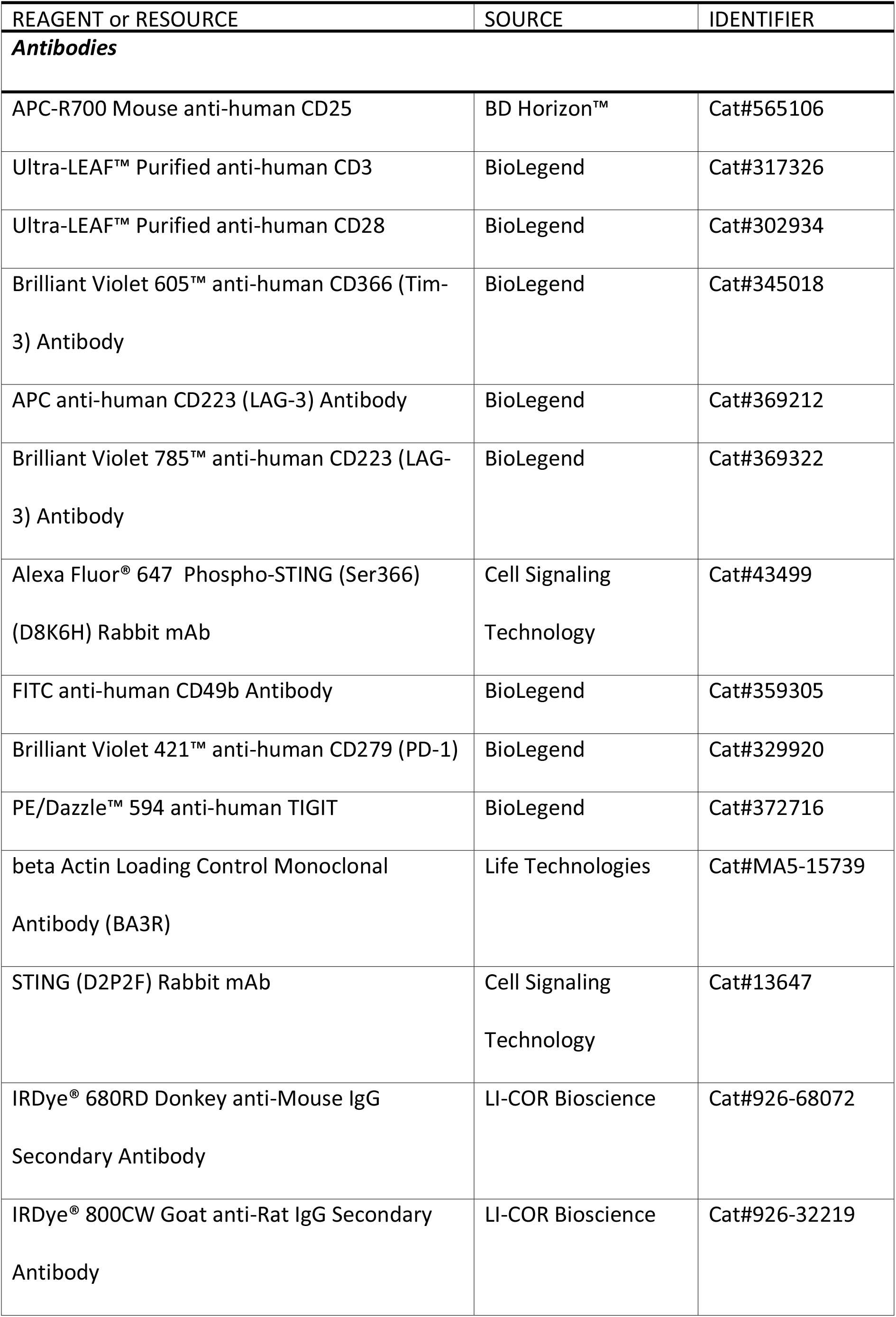
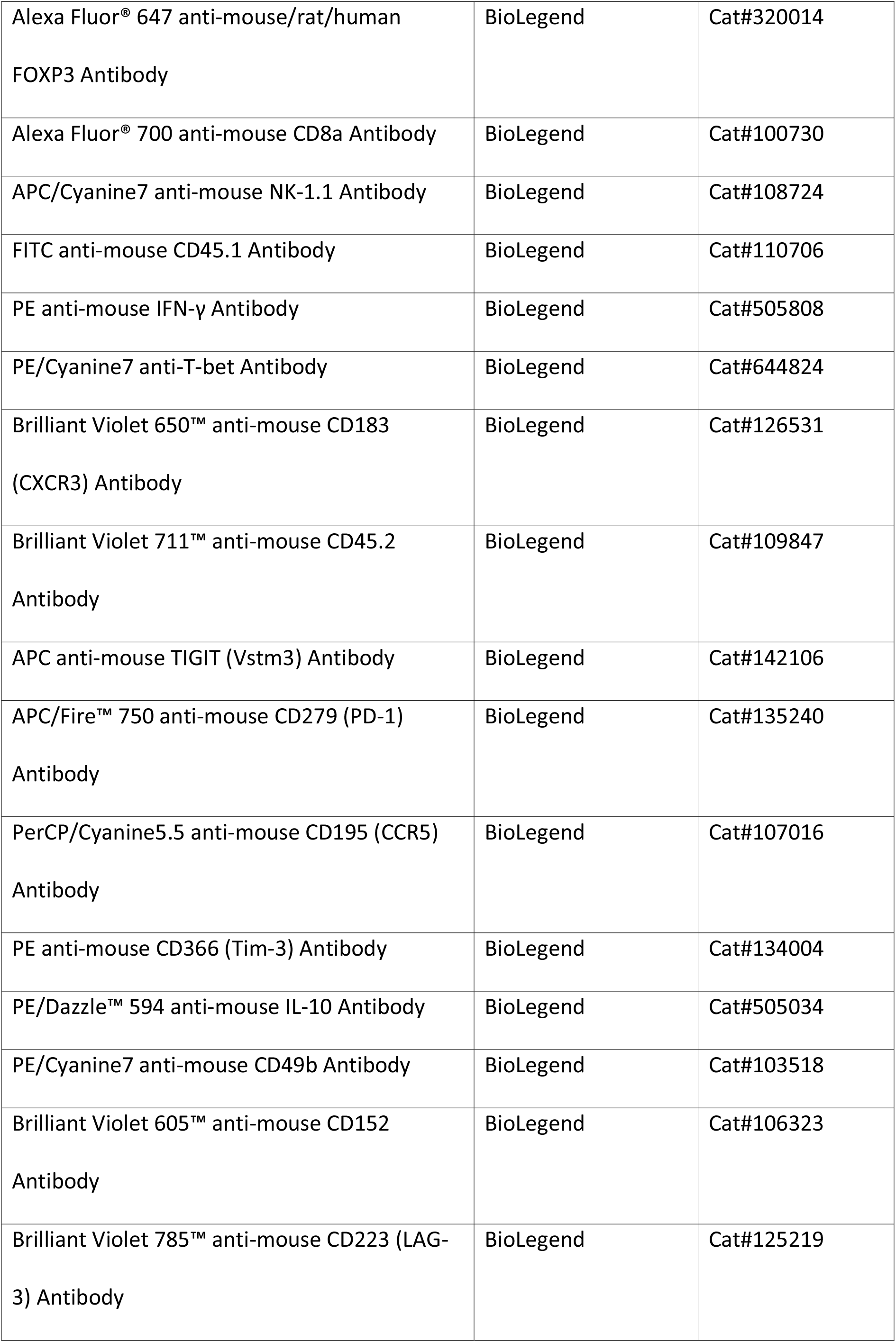
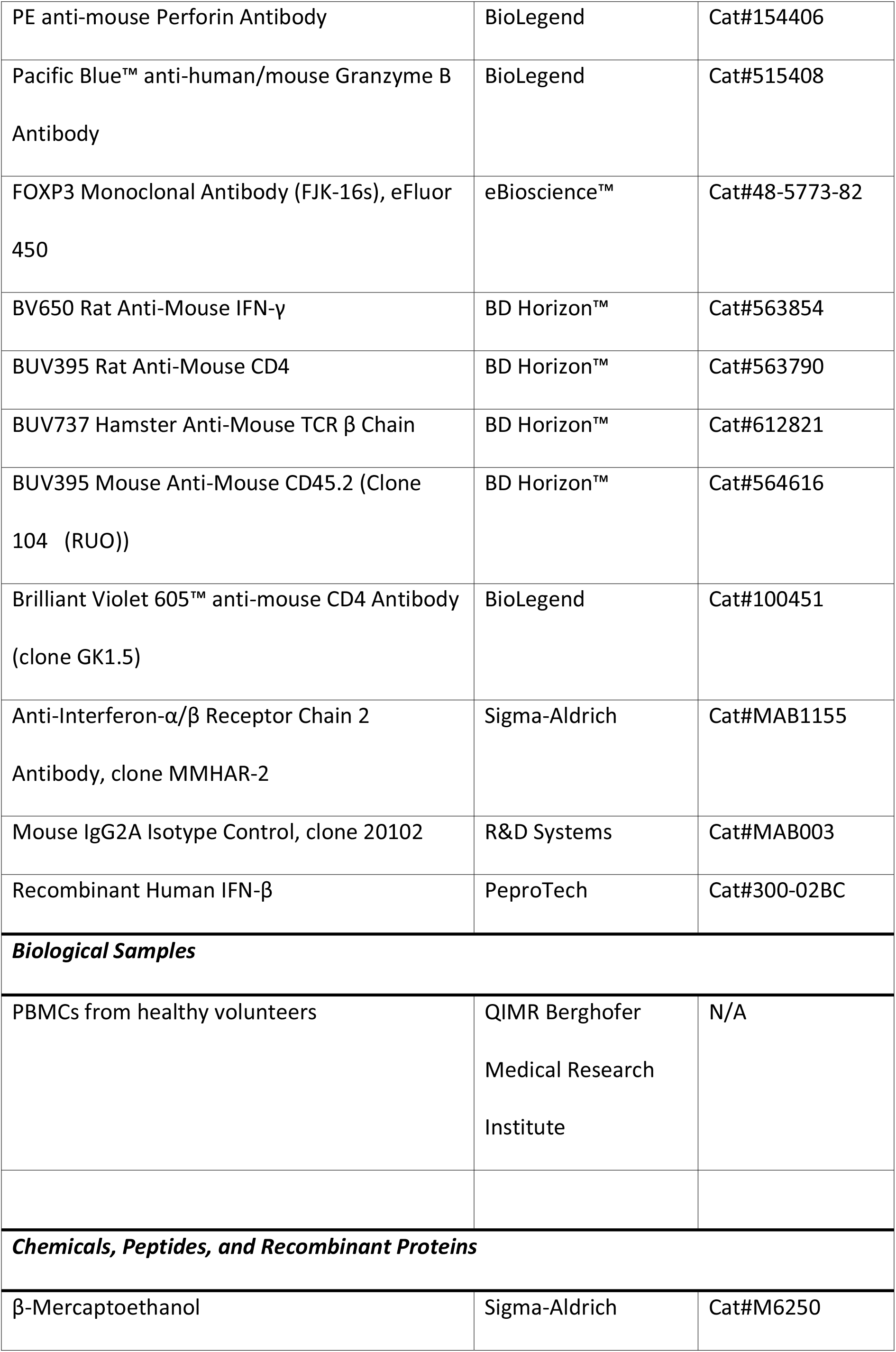
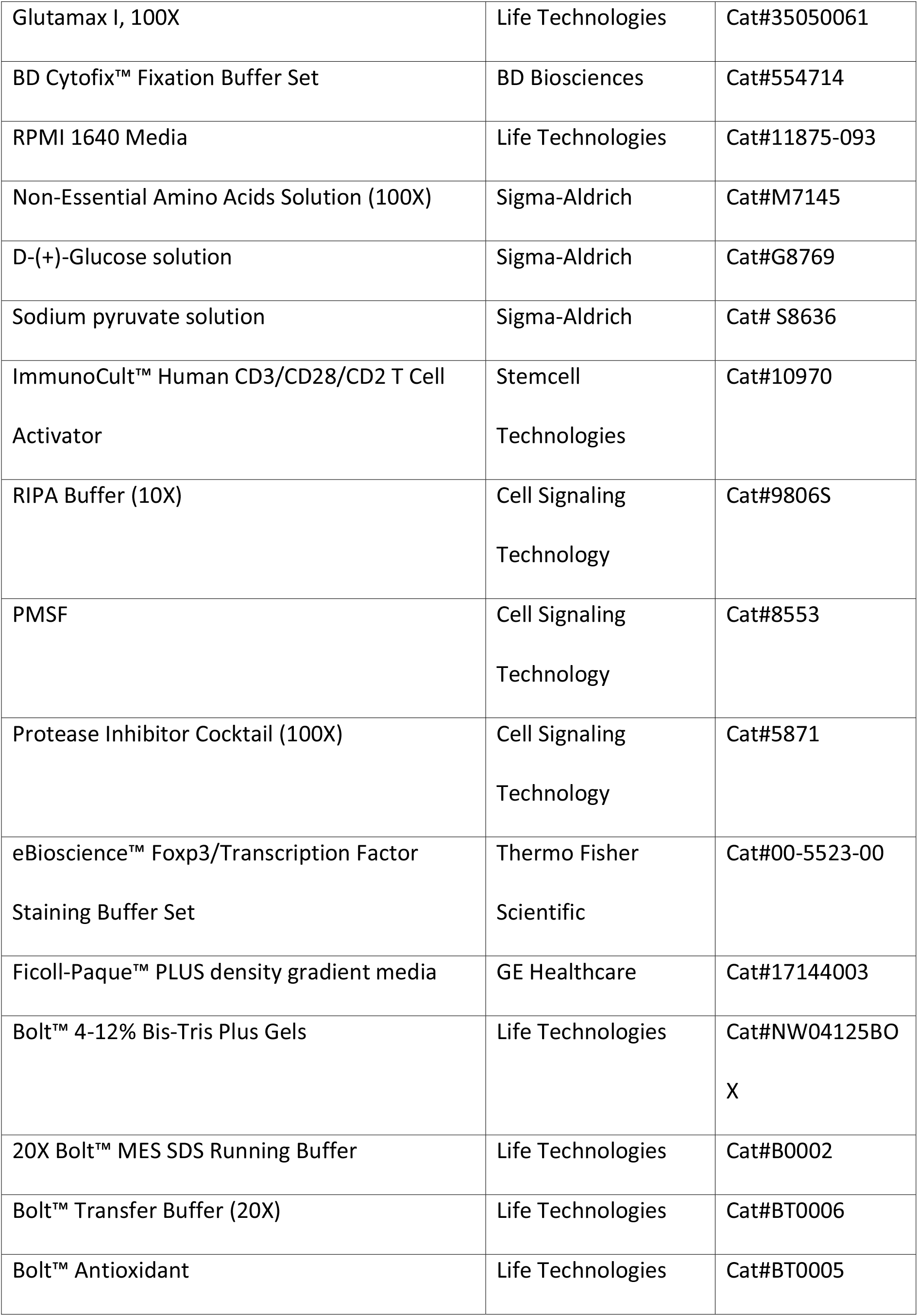
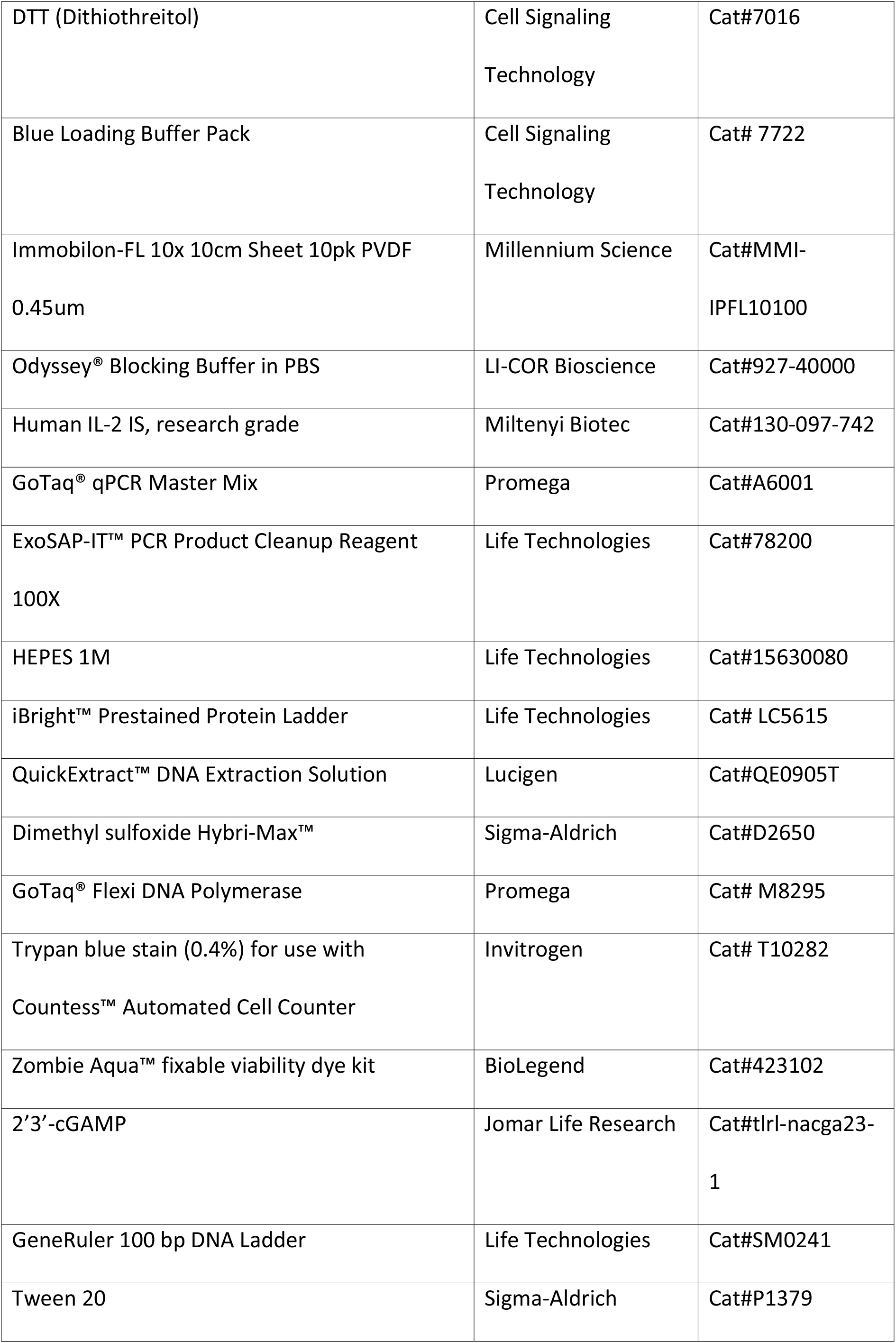
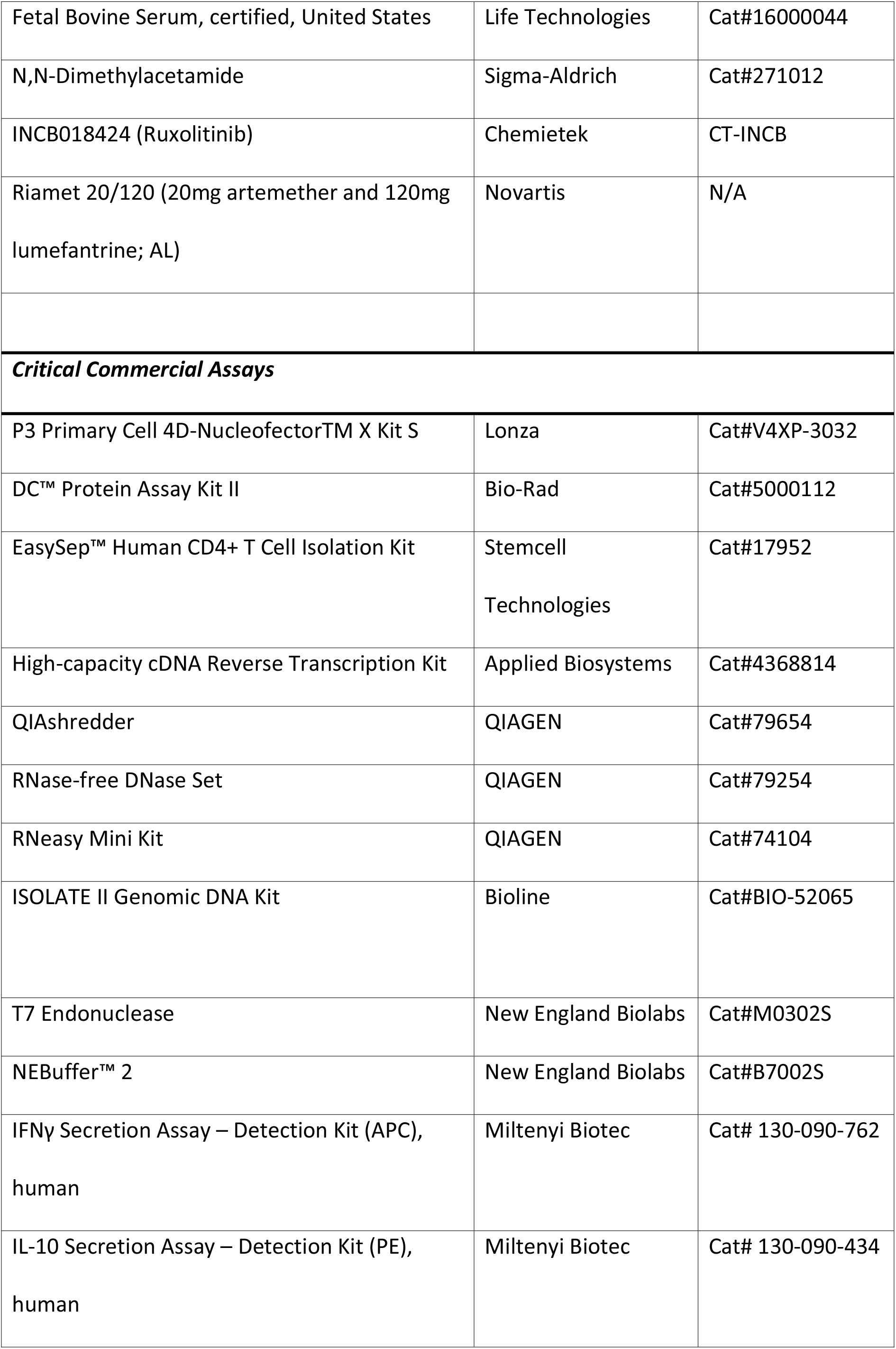
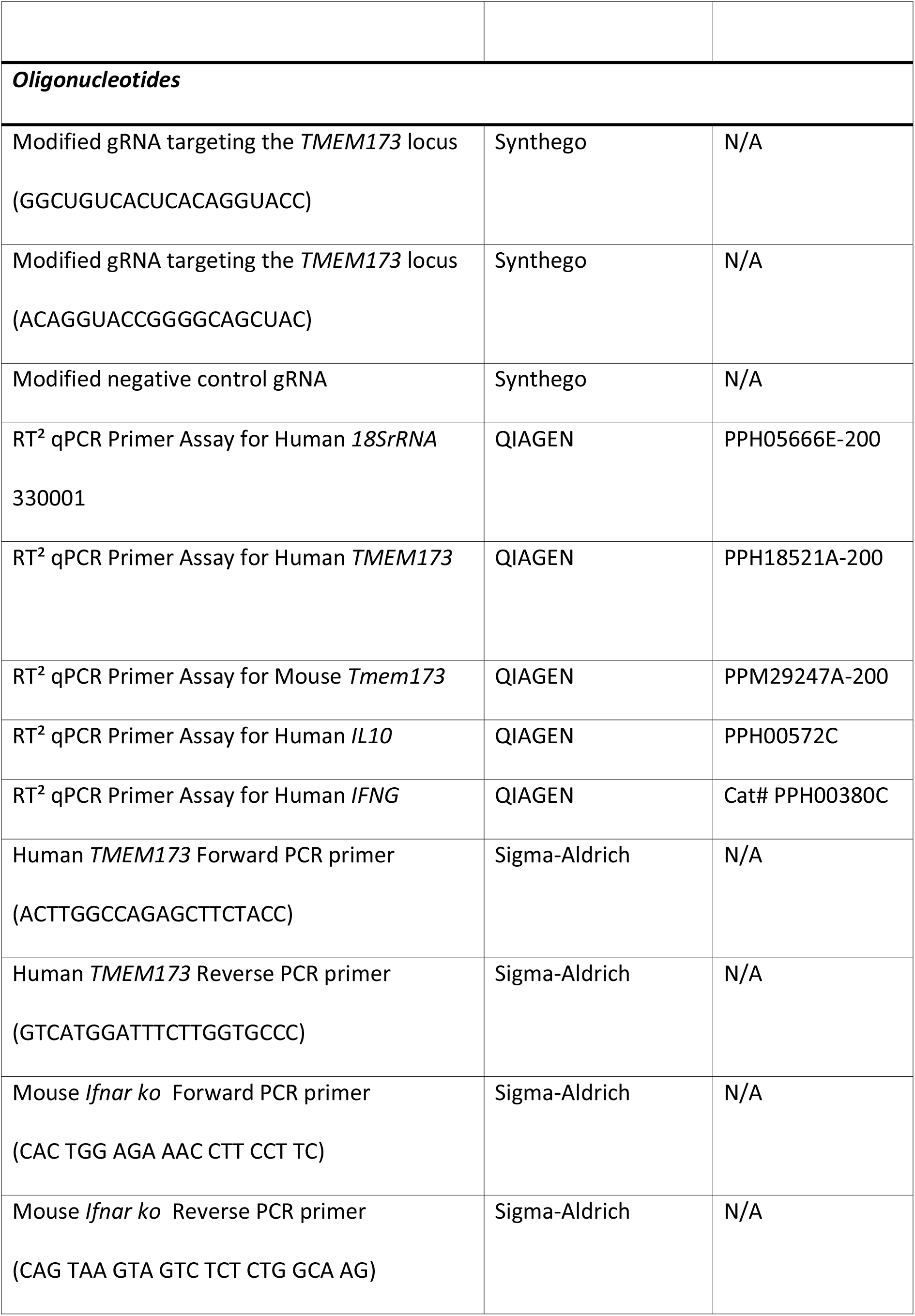
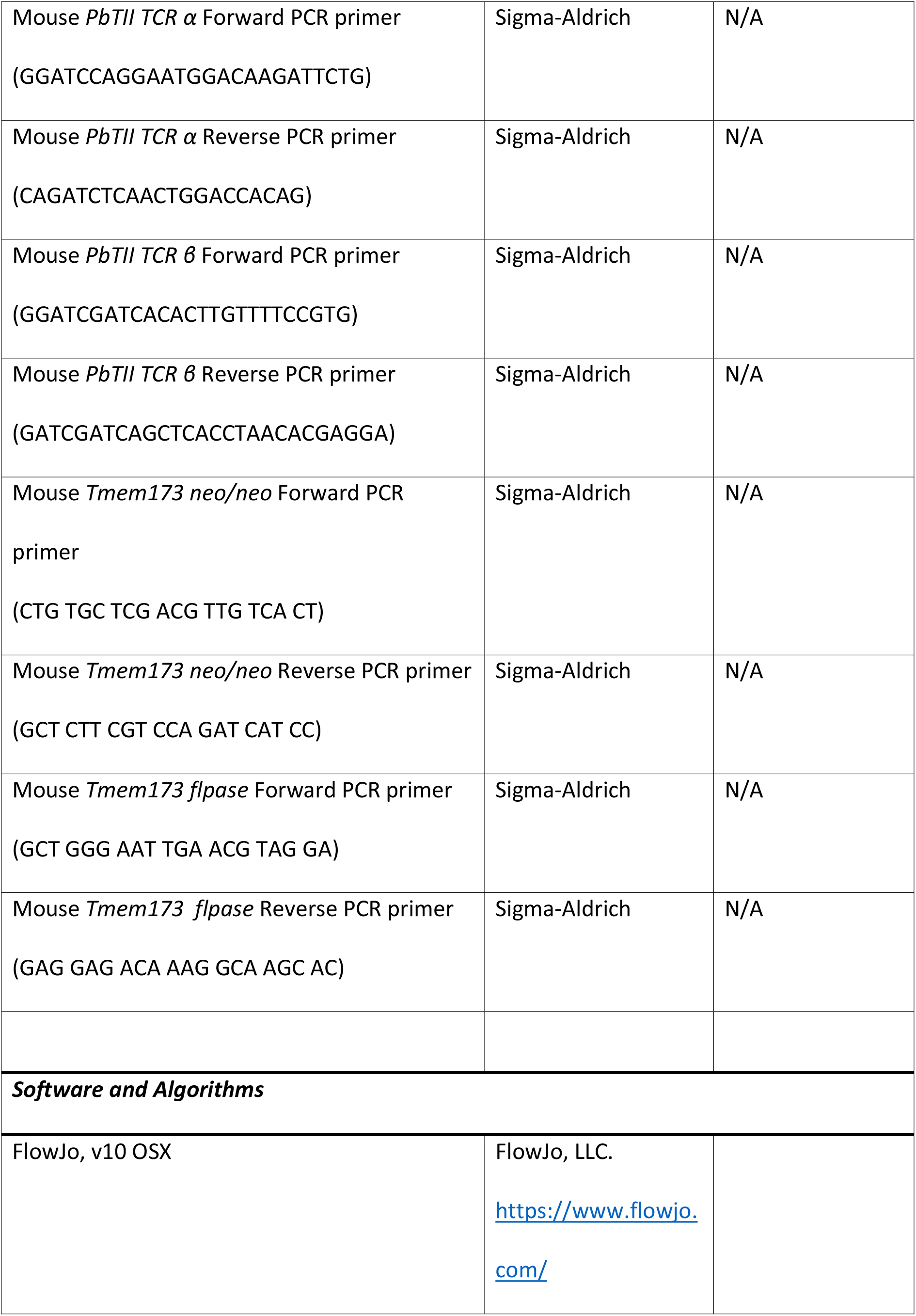
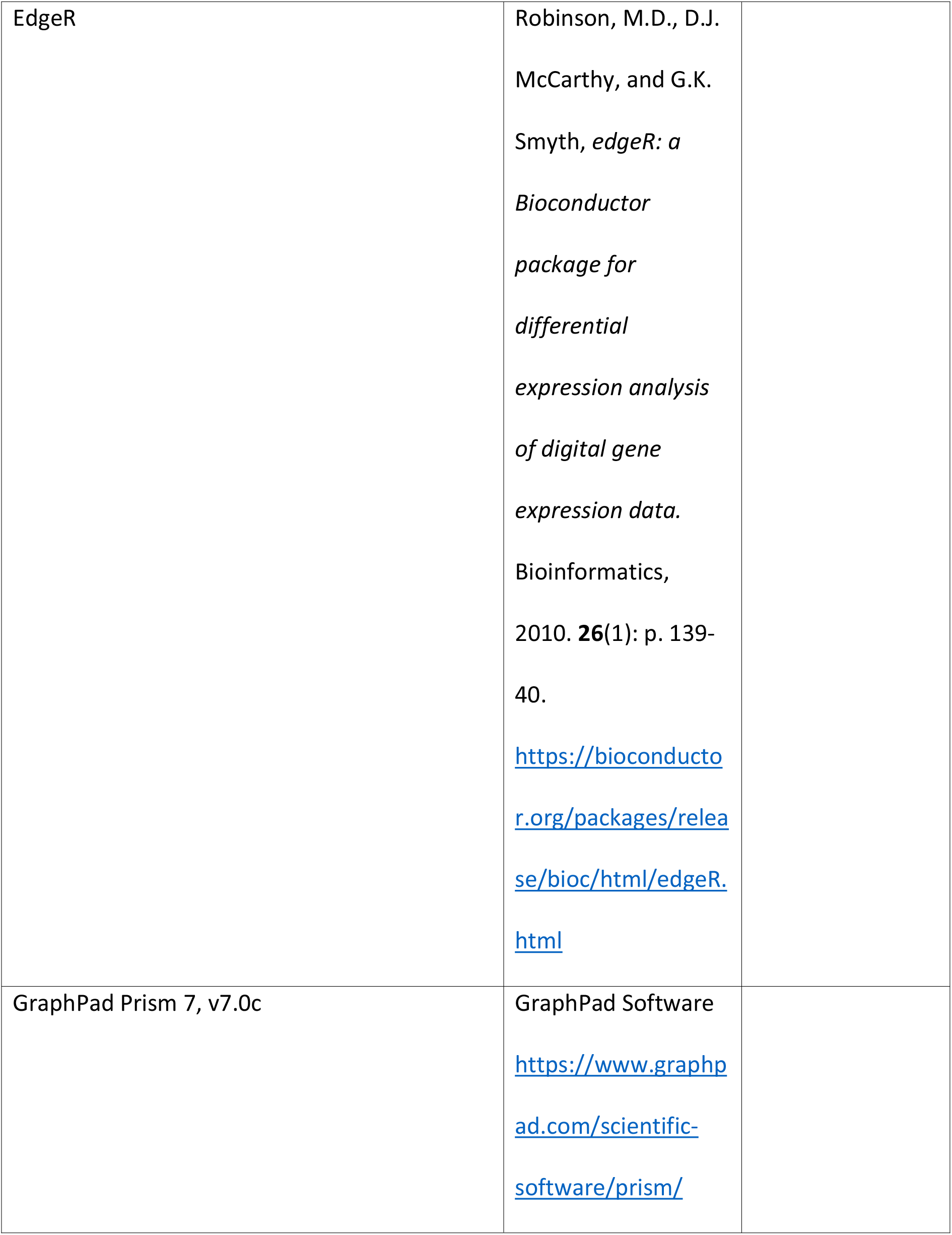
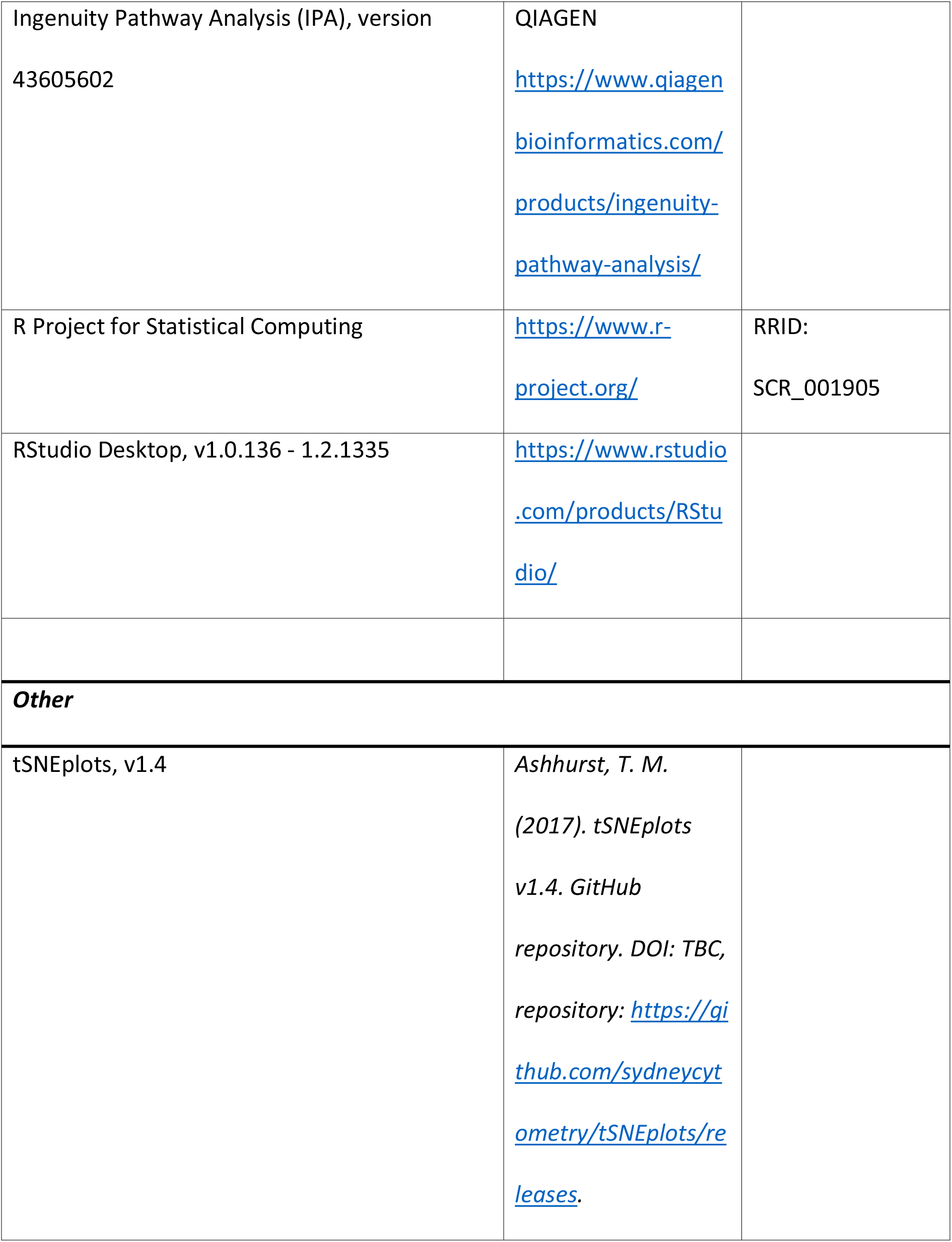
Key resources.

### Western blotting

Cells were lysed with 500 μl RIPA buffer (Cell Signaling Technology, Danvers, MA) with 1X PMSF (Cell Signaling Technology) and protease inhibitor cocktail (Cell Signaling Technology) per 10^7^ cells. According to the manufacturer’s protocol, the protein concentration was measured using a DC Protein Assay Kit II (Bio-Rad). The cell lysate was denatured at 75 °C for 10 minutes, and the protein was separated by Bolt 4-12 % (w/v) Bis-Tris Plus Gels (Life Technologies), transferred onto a PVDF membrane (Millennium Science, Mulgrave, Australia). The membrane was blocked with 50 ml Odyssey Blocking Buffer (LI-COR Bioscience, Lincoln, NE) at room temperature (RT) RT for 1 hour, and then incubated with 1/1000 diluted anti-STING monoclonal antibody (mAb) (Cell Signaling Technology) at 4°C overnight. On the second day, the membrane was washed and incubated with 0.2 μg/ml diluted IRDye 680RD donkey anti-mouse IgG secondary antibody and IRDye 800CW goat anti-rat IgG secondary antibody (LI-COR Bioscience, key resource table) at RT for 30 minutes. The signal was detected using the Licor Odyssey CLx (Millennium Science).

### Cytometric bead array (CBA)

A human Th1/Th2/Th17 Cytokine Kit (BD Biosciences) was used to measure the production of cytokines as per the manufacturer’s instructions. Assays were run on a four laser BD LSRFortessa Cell Analyzer (BD Biosciences). CBA data were analysed using the BD CBA FCAP Array Software v3.0.

### RNA extraction

Cells were washed and lysed with 350 µl RLT buffer (QIAGEN, Hilden, Germany). According to the manufacturer’s instructions, RNA was extracted using the RNeasy Mini Kit (QIAGEN). The concentration of RNA (ng/μl) and sample purity (260/280 ratio) was measured using the NanoDrop 2000 UV-Vis Spectrophotometer (Thermo Fisher Scientific). Extracted RNA was reverse transcribed to complementary DNA (cDNA) using the High-capacity cDNA Reverse Transcription Kit (Applied Biosystems, Waltham, CA), as per manufacturer’s instructions.

### Real-time quantitative polymerase chain reaction (RT-qPCR)

GoTaq qPCR Master Mix (Promega) was used in a final reaction volume of 10μl containing 10 ng of template cDNA. RT-qPCR was performed in Hard-Shell 384-Well Plates, thin wall, skirted, clear/clear (Bio-Rad), sealed with Microseal ‘B’ PCR Plate Sealing Film (Bio-Rad) on the QuantStudio 5 Real-Time PCR System (Applied Biosystems). Relative quantification was performed using the comparative C_T_ method relative to the housekeeping gene *18S* rRNA because it was shown to be a stable housekeeping gene for human T lymphocytes (Bas et al., 2004).

### Bulk RNA sequencing

RNA was extracted from mouse cells using the RNeasy Mini Kit (Qiagen), according to the manufacturer’s instructions. RNase-free DNase Set (Qiagen) was used to remove potential DNA contamination. RNA integrity was accessed using the RNA 6000 Pico Kit (Agilent Technologies, Santa Clara, CA). The cDNA libraries were prepared using NEBNext Single Cell/Low Input RNA Library Prep Kit for Illumina 96 reactions (New England Biolabs). Libraries were quantified using the KAPA Library Quantification Kit (Roche, Indianapolis, IN) and subsequently sequenced on the Illumina NextSeq550 platform (Illumina, San Diego, CA) as a paired-end 75 cycle run (performed by the Sequencing Facility at QIMR). 12.3 – 14.5 million paired reads were obtained per sample.

### RNAseq analysis

Sequence reads were trimmed for adapter sequences using Cutadapt (Martin, 2011) version 1.9 and aligned using STAR (Dobin et al., 2013) version 2.5.2a to the *Mus musculus* GRCm38 assembly with the gene, transcript, and exon features of Ensembl (release 70) gene model. Quality control metrics were computed using RNA-SeQC (DeLuca et al., 2012) version 1.1.8 and expression was estimated using RSEM (Li and Dewey, 2011) version 1.2.30. All downstream RNAseq analysis was performed using R version 3.6.2 (key resource table) and figures were generated using ggplot2 (key resource table). Differential expression analysis was performed using the quasi-likelihood pipeline from edgeR version 3.28.0 (Robinson et al., 2010; Robinson and Oshlack, 2010). Specifically, only protein-coding genes that passed the minimum expression filter using edgeR’s filterByExpr function with default settings were kept for further analysis. The glmQLFit function (Lun et al., 2016) was used to fit a quasi-likelihood negative binomial generalized log-linear model to the read counts for each gene. Using the glmTreat function (McCarthy and Smyth, 2009), we then tested for differential expression between PbTII^Δ*Sting*^ versus PbTII^WT^ cells at day 4 p.i., relative to a minimum fold change threshold of log2(1.5). Differentially expressed genes (DEGs) were determined using a false discovery rate (FDR) < 0.05. Gene set enrichment analysis (GSEA, key resource table) was used to identify classes of differentially expressed genes between experimental and control samples. A heat map was generated by MORPHEUS (key resource table). The scale of the heat map was determined by the differences of LogCPM between paired PbTII^Δ*Sting*^ and PbTII^WT^ cells from the same host.

### Ingenuity pathway analysis (IPA)

Gene ID, log_2_ fold-change (logFC) of gene expression, and adjusted *P*-value were used to input into IPA (version 43605602; QIAGEN). Initial interrogation of each dataset was performed using default values and parameters set on IPA. Upstream pathway analysis was performed on *IL10*. The tool *Grow* was used, direct and indirect interactions were selected, and the molecules/gene interactions related to the RNAseq data were identified.

## Supplemental information

**Figure S1.**
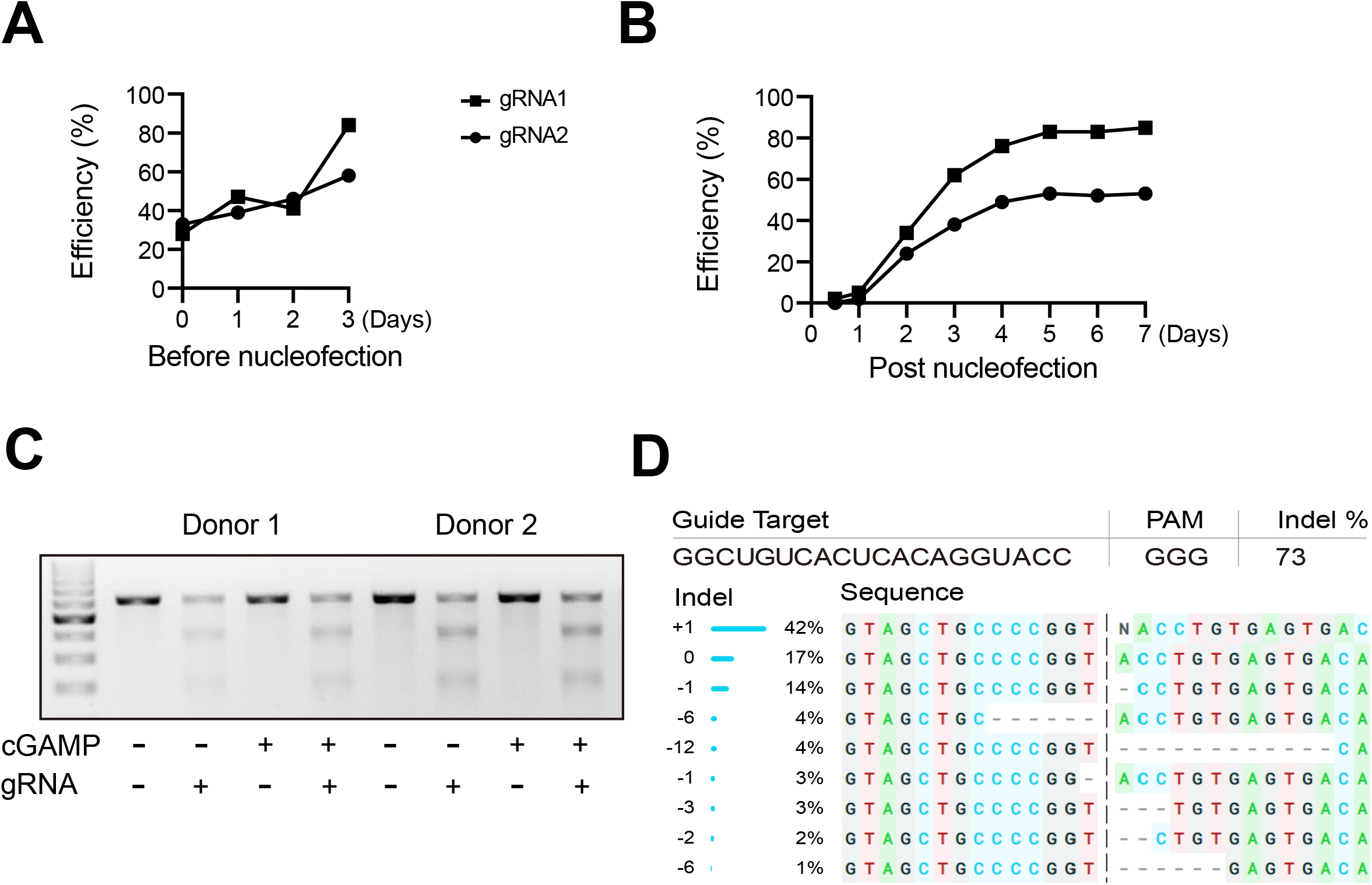
Optimization of the CRISPR/Cas9 gene editing protocol for human activated CD4^+^ T cells. **A**, Peripheral blood CD4^+^ T cells from a healthy volunteer were stimulated with αCD3ε and αCD28 mAbs plus IL-2 for 0, 1, 2 and 3 days before nucleofection with two different *TMEM173* guide RNAs (gRNA), and were stimulated for another 3 days, as above, before analysis. Gene-editing efficiency was measured by Big Dye Sequencing. **B**, CD4^+^ T cells from healthy volunteers were stimulated with αCD3ε and αCD28 mAbs plus IL-2 for 3 days before nucleofection with two different *TMEM173* gRNA and were then stimulated for 0 to 7 days before Big Dye Sequencing to calculate the efficiency of gene editing. **C**, CD4^+^ T cells were stimulated for 3 days, as above, transfected with *TEMEM 173* gRNA1, then stimulated for another 3 days with the addition of cGAMP 18 hours before analysis, as indicated. Representative T7 endonuclease I mismatch assay showing *TMEM173* modifications. **D**, Representative Inference of CRISPR Edits (ICE) analysis (https://ice.synthego.com/#/) of Big Dye Sequencing data showing the editing efficiency, deletions or insertion of single nucleotides or fragments (Indels).

**Figure S2.**
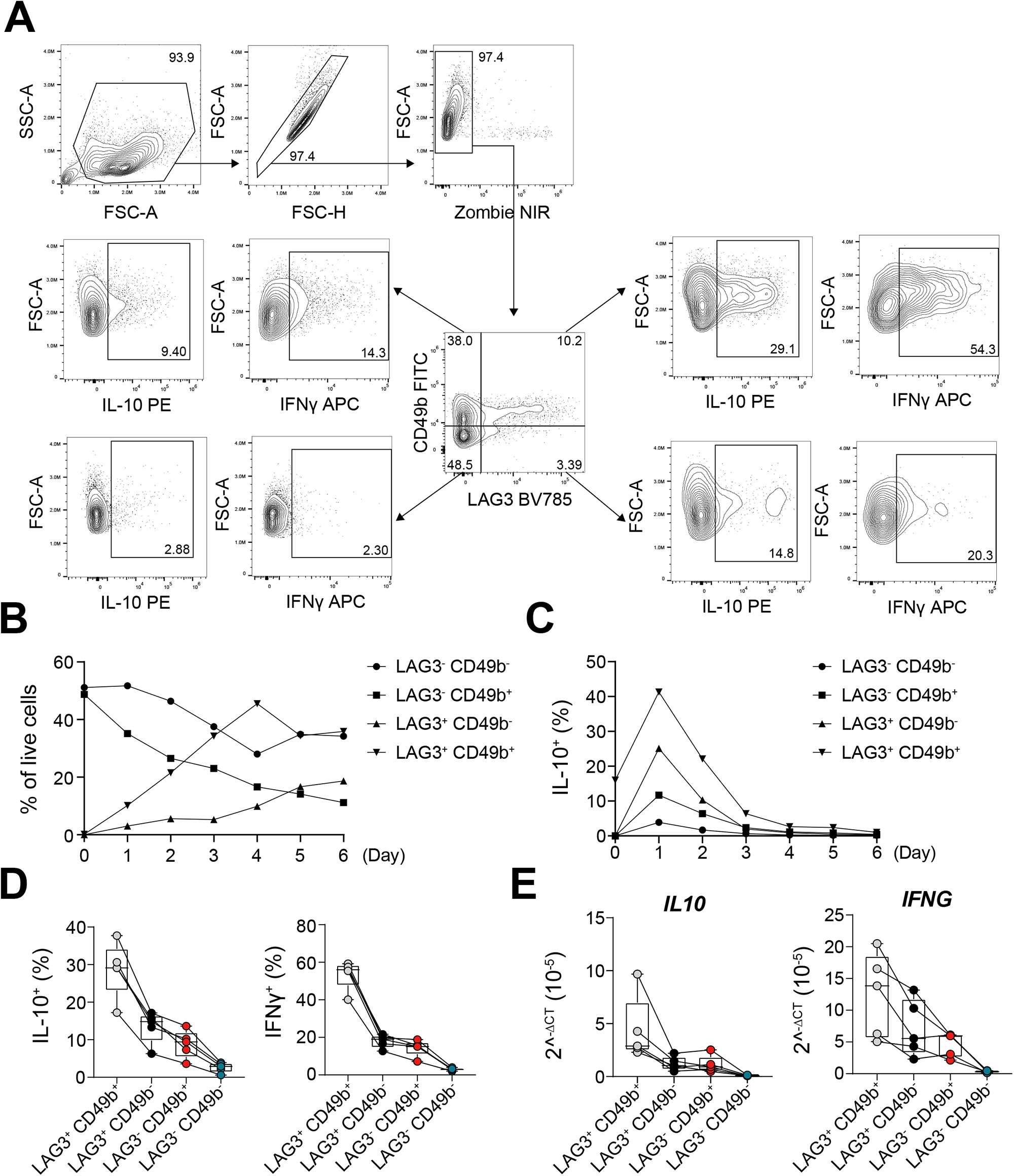
LAG3 and CD49b are surrogate markers for Tr1 cells. **A**, Peripheral blood CD4^+^ T cells from healthy volunteers (n = 4) were stimulated with αCD3ε and αCD28 mAbs plus IL-2 for 0 to 6 days. LAG3, CD49b, IL-10 and IFNγ expression was measured every day. The gating strategy for IL-10^+^, IFNγ^+^, and LAG3^+^ CD49b^+^ CD4^+^ T cells is shown. **B**, Changes in the frequency of LAG3^+^ CD49b^+^ CD4^+^ T cells from day 0 to day 6. **C**, The kinetics of IL-10 expression by CD4^+^ T cell subsets based on LAG3 and CD49b expression. **D**, Purified CD4^+^ T cells from healthy volunteers were stimulated with αCD3ε and αCD28 mAbs plus IL-2 for 1 day. Half of the cells were stained for LAG3, CD49b, IL-10 and IFNγ, while the other half were sorted into CD4^+^ T cell subsets based on LAG3 and CD49b expression. The expression of IL-10 and IFNγ protein by cell subsets is shown. **E**, *IL10* and *IFNG* mRNA in sorted LAG3 and CD49b CD4^+^ T cell subsets was measured by qPCR. Data was normalized to the housekeeping gene *18S* rRNA.

**Figure S3.**
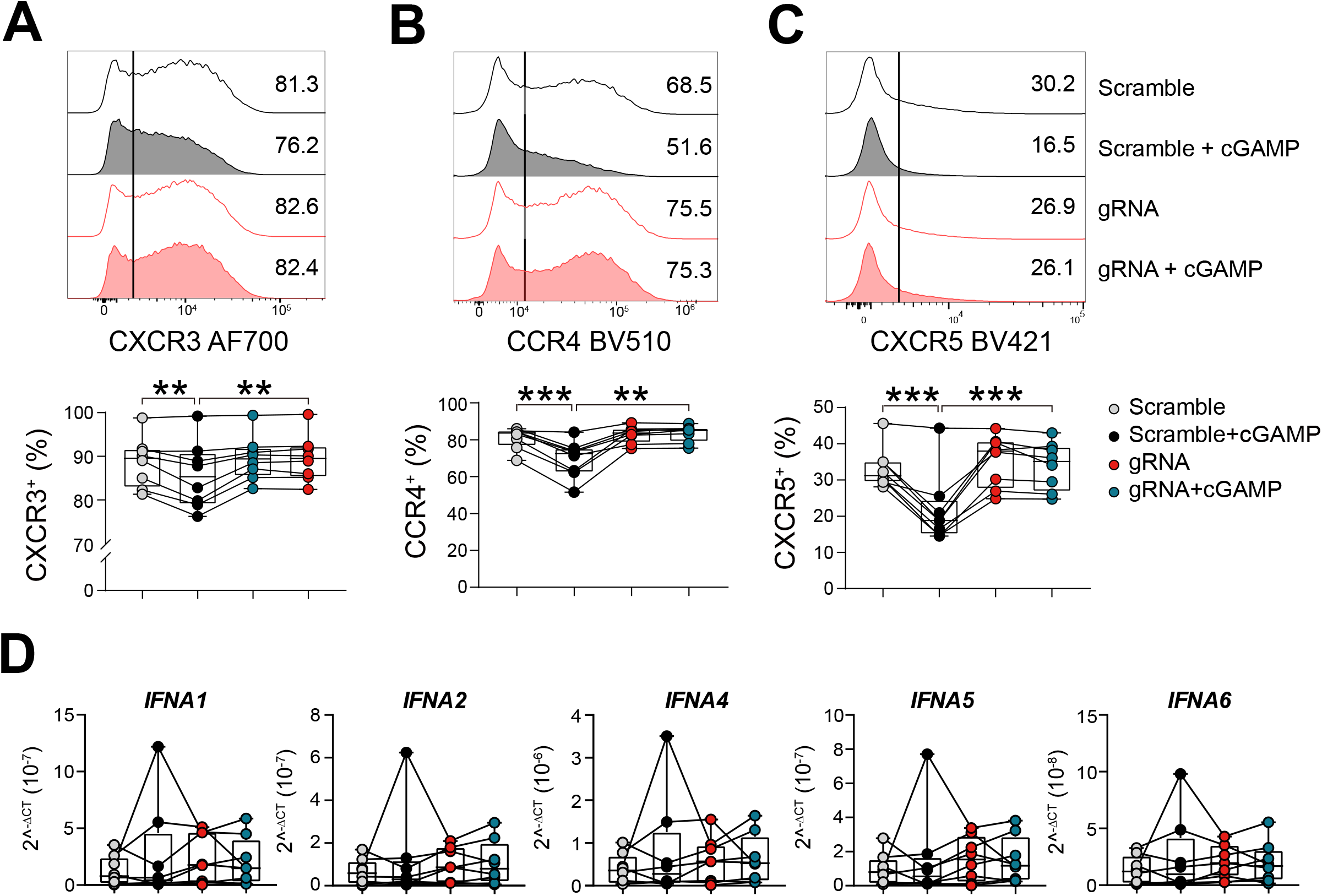
STING inhibits the expression of CXCR3^+^, CCR4^+^ and CXCR5^+^ by CD4^+^ T cells. **A**, **B**, **C**, Representative histograms and enumeration showing the frequency of CXCR3^+^, CCR4^+^ and CXCR5^+^ CD4^+^ T cells, respectively, following CRISPR/Cas9 to modify *TMEM173* expression. n=8, paired t-test. **D**, *IFNA1*, *IFNA2*, *IFNA4*, *IFNA5* and *IFNA6* mRNA was measured by qPCR. Data was normalized to the housekeeping gene *18S* rRNA and was Log2-transformed for statistical analysis. n=8, paired t-test. Comparisons between scramble vs scramble + cGAMP, scramble + cGAMP vs gRNA + cGAMP, gRNA + cGAMP vs gRNA were made. **P < 0.01, ***P < 0.001.

**Figure S4.**
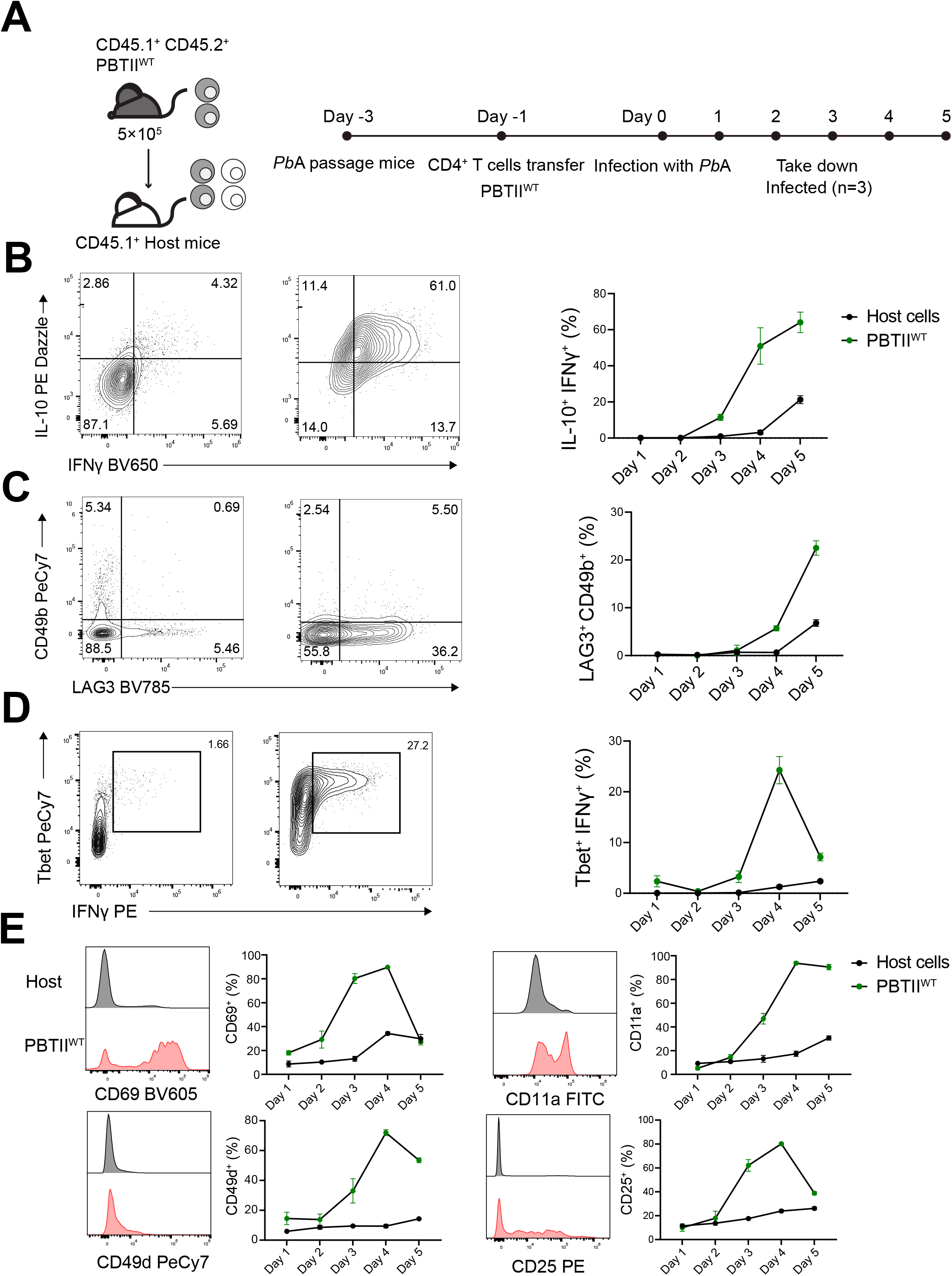
The kinetics of immune PbTII cell expansion in response to *Plasmodium berghei* ANKA (*Pb*A) infection. **A**, 5×10^5^ CD45.1^+^ CD45.2^+^ PbTII^WT^ cells were transferred into the *ptprca* (CD45.1^+^) host mice at day -1. The mice were infected with *Pb*A on day 0 and were taken down from day 1 to day 5 post-infection (p.i.). **B, C, D,** Representative plots and enumeration showing the frequencies of IL-10^+^ IFNγ^+^, LAG3^+^ CD49b^+^ cells and Tbet^+^ IFNγ^+^ PbTII-^WT^ cells from day 1 to 5 p.i.. **E**, Representative histograms and enumeration showing the expression of the activation markers CD69, CD11a, CD49d and CD25 by PbTII^WT^ cells from day 1 to 5 p.i..

**Figure S5.**
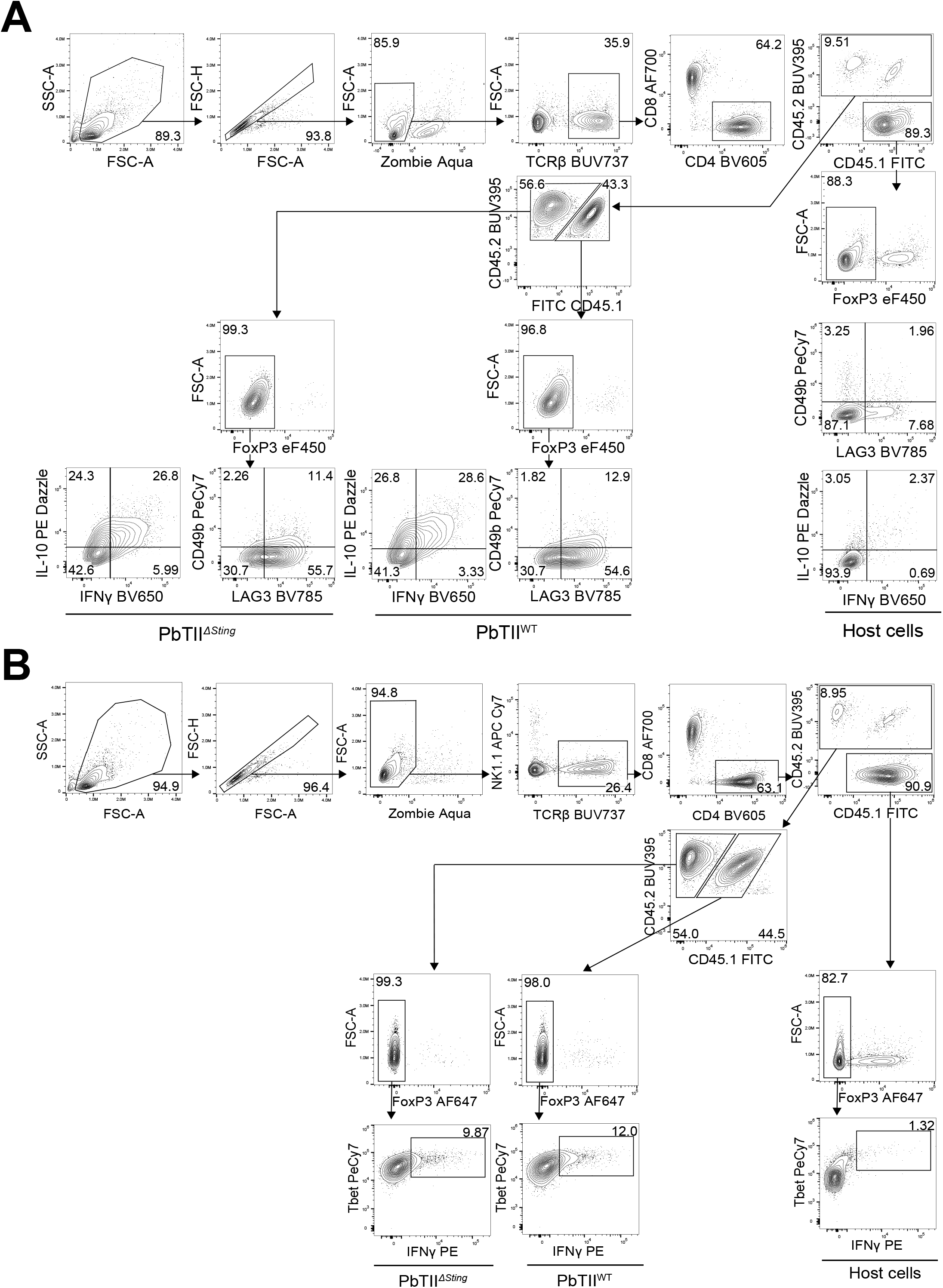
Identification of Tr1 and Th1 PbTII^Δ*Sting*^ and PbTII^WT^ cells. Live cells were selected, followe by gating on CD4^+^ T cells. These cells were then divided into adoptively transferred PbTII^Δ*Sting*^ and PbTII^WT^ cells, based on congenic marker expression. **A**, IL-10^+^ IFNγ^+^ and LAG3^+^ CD49d^+^ PbTII^Δ*Sting*^ and PbTII^WT^ Tr1 cells, and **B**, Tbet^+^ IFN γ^+^ PbTII^Δ*Sting*^ and PbTII^WT^ Th1 cells were then assessed.

**Figure S6.**
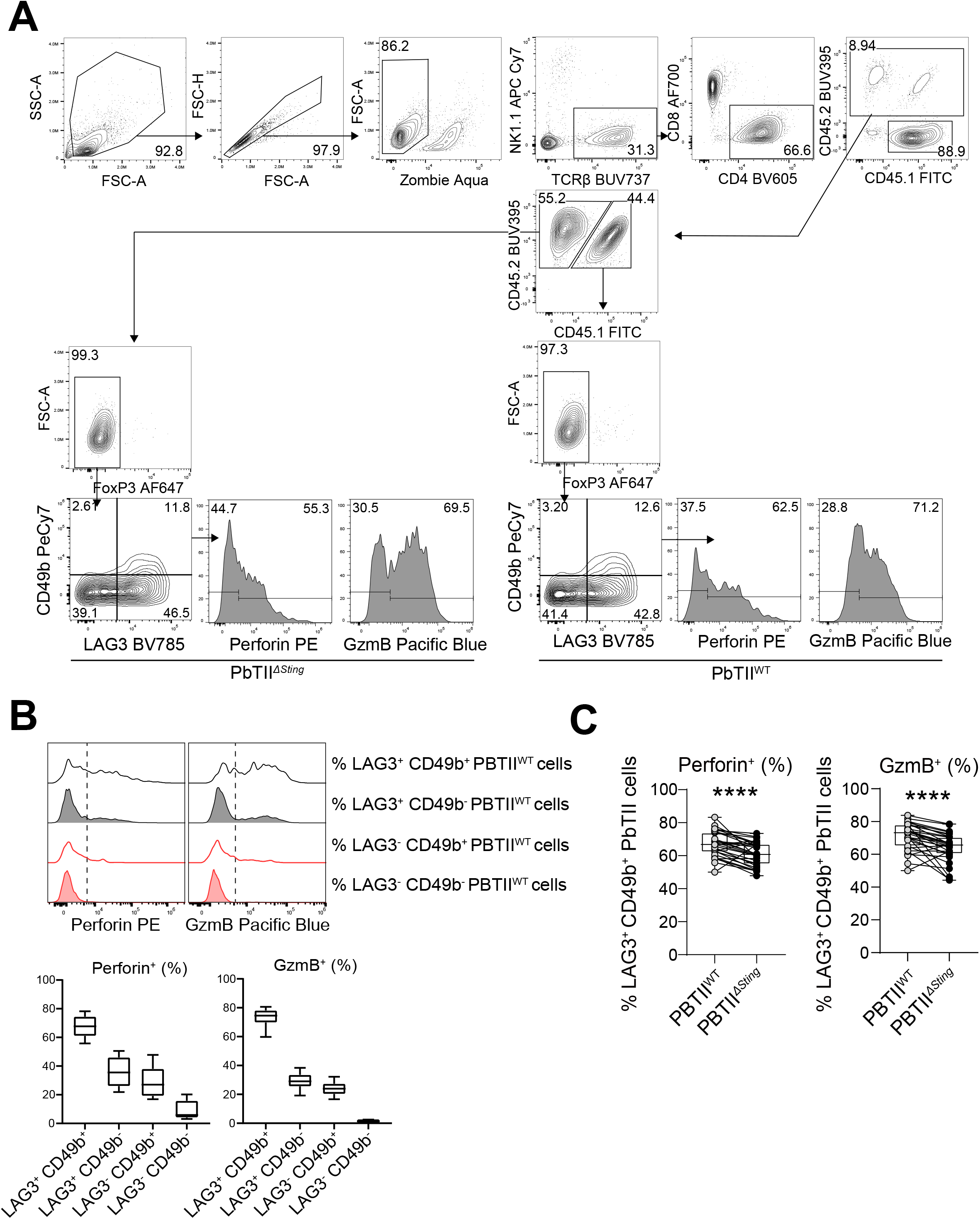
STING promotes the expression of cytotoxic molecules by Tr1 cells in experimental malaria. Spleen cells from day 4 post-infection with *Plasmodium berghei* ANKA were stimulated with monensin, PMA and ionomycin for 3 hours *ex vivo* before antibody staining. **A**, The gating strategy used to assess the frequency of Tr1 cells and their expression of cytotoxic molecules. **B**, Representative histograms and enumeration showing the expression of perforin and granzyme B (GzmB) by PbTII^WT^ cell subsets based on LAG3 and CD49b expression. **C**, The enumeration showing the frequency of perforin^+^ or GzmB^+^ PbTII^Δ*Sting*^ and PbTII^WT^ Tr1 cells. n = 15/group, pooled from 3 independent experiments, paired t test. ****P < 0.0001.

**Figure S7.**
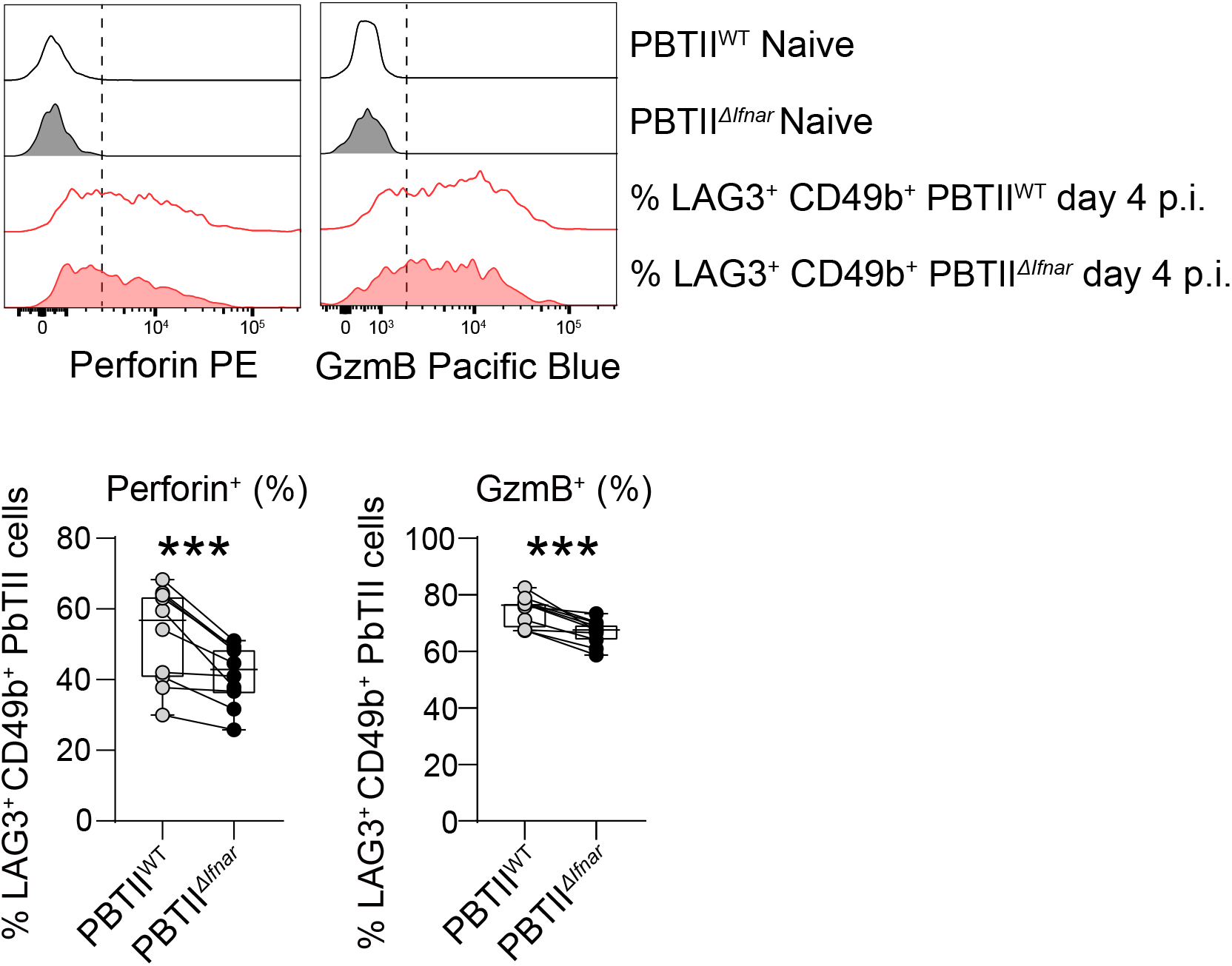
Type I IFN signalling promotes the expression of cytotoxic molecules by Tr1 PbTII cells during malaria. Spleen cells from day 4 post-infection with *Plasmodium berghei* ANKA were stimulated with monensin, PMA and ionomycin for 3 hours *ex vivo* before antibody staining. Representative histogram and enumeration showing the frequencies of perforin^+^ or granzyme B (GzmB)^+^ Tr1 cells in PbTII^Δ*Ifnar*^ and PbTII^WT^ cells. n = 10/group, pooled from 2 independent experiments, paired t test. ***P < 0.001.

**Figure S8.**
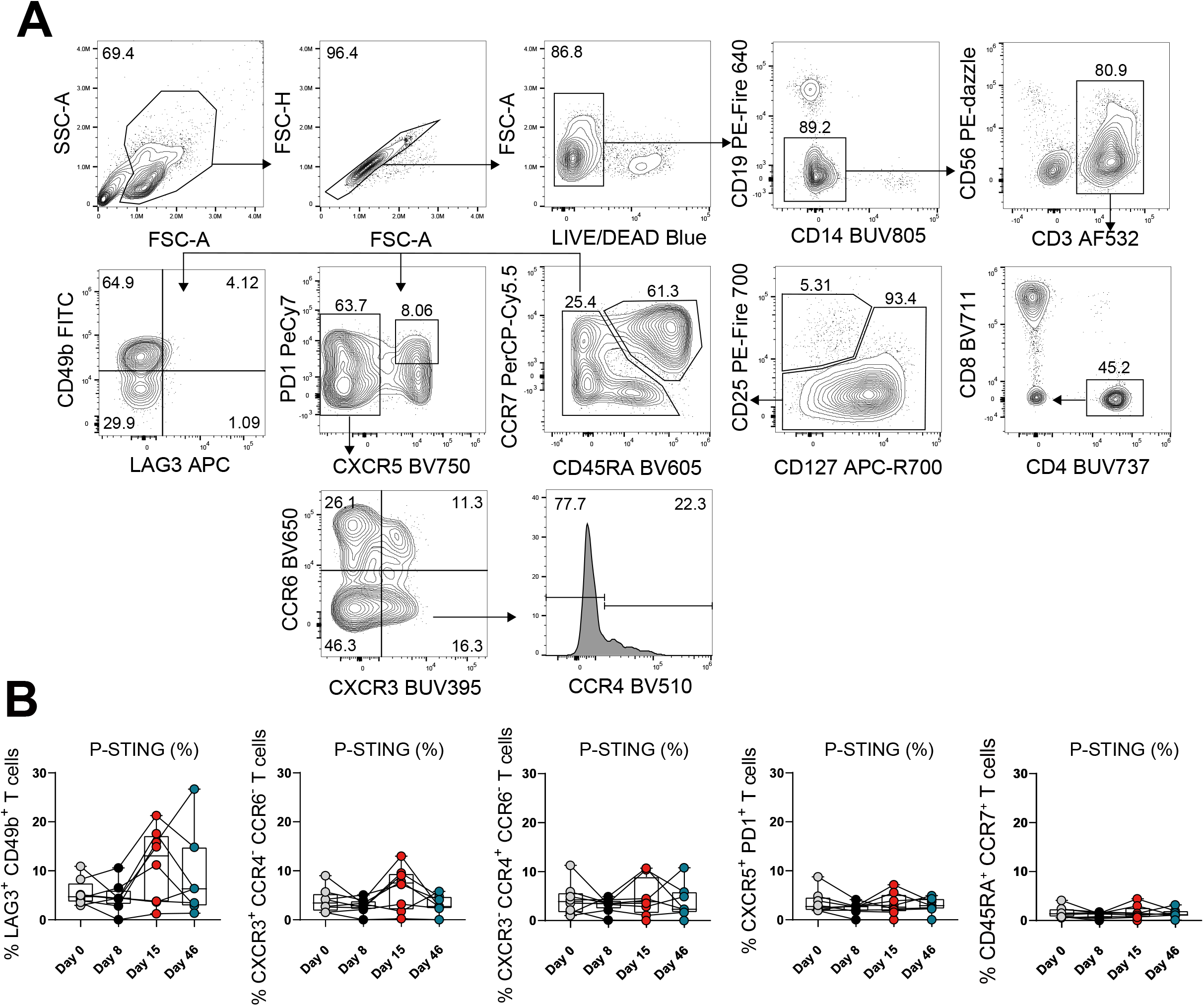
Tr1 cells from humans infected with *Plasmodium falciparum* are more sensitive to STING activation. **A,** The gating strategy used to identify CD4^+^ T cell subsets. B, The frequency of different Th cell subsets that contained phosphorylated STING (P-STING) from day 0 post-infection (p.i.) to 46 p.i. in the presence of cGAMP for 18h.

**Figure S9.**
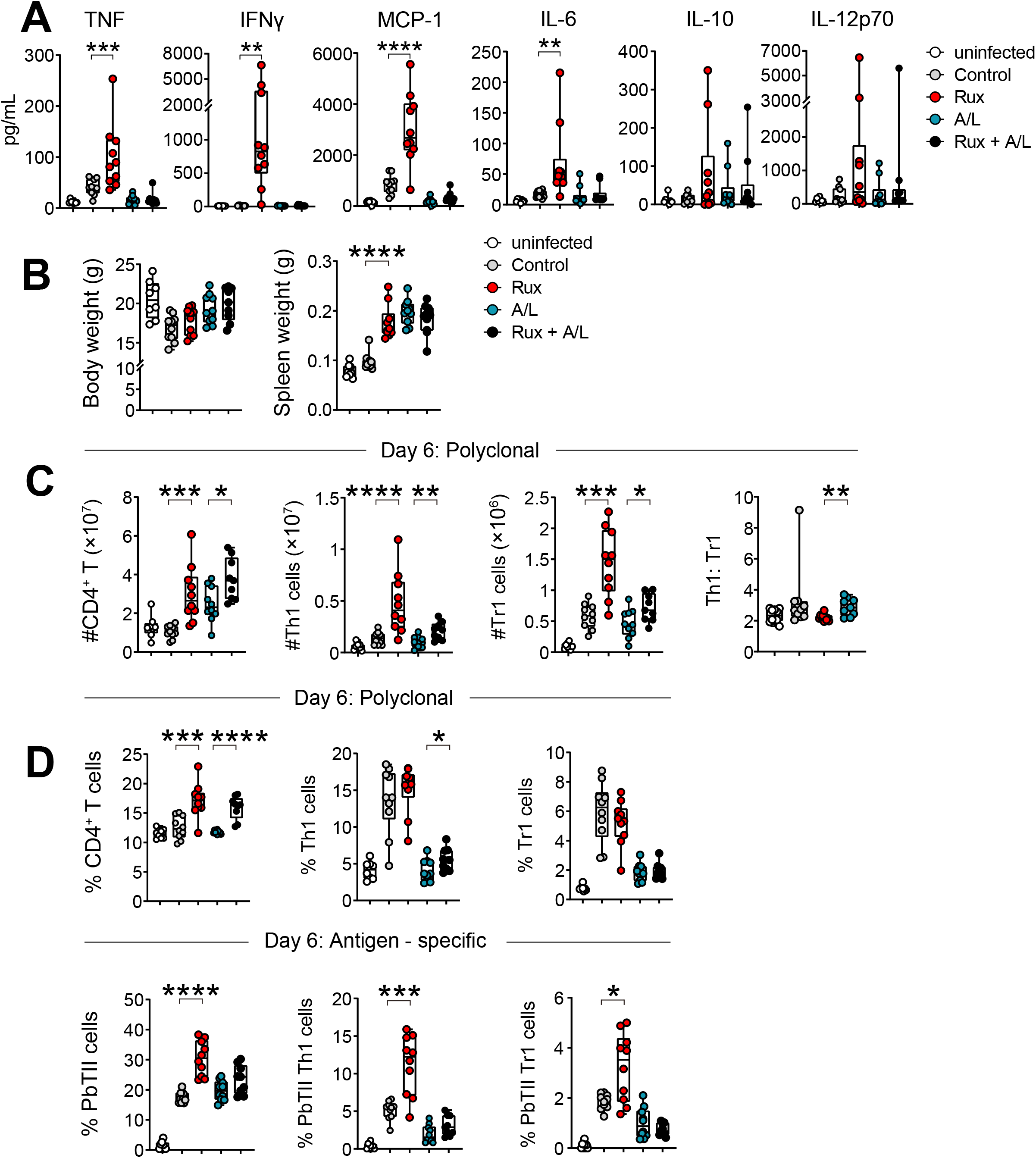
Ruxolitinib treatment improves anti-parasitic CD4^+^ T cell responses in a lethal malaria model with *Plasmodium berghei* ANKA. C57BL/6J mice were infected with 1×10^5^ luciferase transgenic *Plasmodium berghei* ANKA (PbA-luc) pRBCs intravenously (i.v.) and treated with either DMAC (control), 60mg/kg of Ruxolitinib (Rux), medium dose of artemether/lumefantrine (A/L; 1.5mg/kg of artemether combined with 9mg/kg of lumefantrine per mouse) or Rux combined with A/L beginning at day 3 post-infection (p.i.) until day 5 p.i., twice daily and then assessed on day 6 p.i.. **A**, Serum cytokine levels, as well as **B**, body and spleen weights were measured at day 6 p.i.. **C**, Total CD4^+^ T cell, Th1 cell and Th1 cell numbers in the spleen at day 6 p.i. are shown for each group, as well as Th1:Tr1 cell ratios. **D**, Frequencies of polyclonal and antigen-specific PbTII CD4^+^ T cells, Th1 cells and Tr1 cell in the spleen at day 6 p.i., as indicated, are shown for each group. Boxes show the extent of the lower and upper quartiles plus the median, while the whis-kers indicate the minimum and maximum data points, n= 8-10 mice/group, data are pooled from two representative experiments, *P < 0.05, **P < 0.01, ***P < 0.001, ****P < 0.0001. A, B, C: Mann-Whitney U test.

